# Facilitating NMR Resonance Assignment with Metabolic Tampering

**DOI:** 10.64898/2026.04.29.721603

**Authors:** Danica S. Cui, Evan O. Anderson, Erik Zavala, George P. Lisi, J. Patrick Loria

## Abstract

The ability to assign amino acid resonances in multidimensional NMR spectra of biomolecules is necessary for detailed studies of protein structure and dynamics. Despite creative advances in isotopic labeling, unlabeling and multidimensional NMR experiments, resonance assignment remains a bottleneck in studies of large proteins. In this work, we show that the metabolic flux through biosynthetic pathways of amino acid production during protein expression can be modulated to aid in the identification of resonances in two-dimensional NMR spectra. This straightforward method involves doping ^15^N-enriched minimal media with small amounts of rich natural abundance media to generate unique peak intensity attenuation patterns, producing type-specific signatures of amino acids in two-dimensional ^15^N HSQC experiments. Using three model proteins, IGPS (51 kDa heterodimer), PTP1B (35 kDa), PHPT1 (14 kDa), we show that this method can disentangle several amino acid types, is robust to different expression conditions, and is a useful supplement for triple resonance experiments in protein backbone resonance assignments.

## Introduction

The improvement of NMR technology and methodology has been integral to the advancement of the field and dictates the bounds to which one can push experimental pursuits. Innovations in isotopic labeling^1–5^ pulse sequences ^6–14^, and increases in static magnetic field strengths have made structural and dynamic investigations of large proteins and protein complexes readily achievable ^10,15–20^. The atomic resolution insight afforded by protein NMR spectroscopy obviously requires site-specific assignment of NMR resonances. As such, advances in NMR resonance assignment have proceeded in lock-step with advances in NMR pulse sequences and subsequent mechanistic insight that is obtained for ever larger biomolecules. The expression of proteins in bacterial auxotrophs supplemented with appropriate spin-1/2 labeled amino acids has been utilized to provide resonance assignment of critical amino acid residues and for simplifying the resulting spectra both for NMR and EPR spectroscopy^21–24^.

Related efforts have also sought to simplify NMR spectral assignments through bacterial protein expression in deuterated water supplemented with desired protonated amino acids^25,26^. Exploitation of the glucose metabolic pathways have been further used to aid in the stereospecific assignment of valine and leucine residues^27,28.^

Isotope labeling approaches have also been cleverly combined with heteronuclear filters to simplify the resulting NMR spectra ^29^. It has previously been demonstrated that ^13^C’ resonances in proteins can be assigned through correlation of ^2^H isotope effects on chemical shifts and amide proton exchange rates^30^. Moreover, the use of combinatorial patterns of selective amino acid labeling has aided amino acid type identification in simple two-dimensional HSQC experiments ^31^. However, these detailed NMR studies still rely upon incremental progress in the most basic aspects of the field. Resonance assignment, for example, is critical for atomistic pictures of biomolecules. To further aid in the assignment process a number of ‘cell-free’ protein synthesis methods have been devised that incorporate various combinatorial isotope labeling schemes^32^. The advent of triple-resonance NMR methods offered a breakthrough in the resonance assignment problem and has enabled enormous functional insight into biomolecules ^33–35^. Although, more recently, numerous procedures have sought to simplify, accelerate, and automate the process ^36–41^, however resonance assignments can still represent a substantial bottleneck to further study particularly in larger proteins and protein complexes. Methods that can aid in the unambiguous identification of amino acids, especially in such large systems with complex spectra, are desirable. Here, we present a method that complements some of the recently reported biomolecular assignment techniques, most notably selective unlabeling ^42–44^, which leverages the differential metabolic rates of amino acids to quickly and faithfully identify amino acid type in ^1^H-^15^N heteronuclear single-quantum coherence (HSQC) spectra of the amide backbone. This method of ‘metabolic tampering’ (Fig 1) involves very few sample preparations and maintains its utility in three model proteins that differ in molecular weight, structure, and protein expression conditions. This approach is applicable to standard isotopic labeling protocols and is achieved by doping bacterial growths carried out in spin-1/2 isotope labeled M9 minimal medium, the standard for NMR sample preparation, with small amounts of natural abundance lysogeny broth (LB, 1 – 10%) and subsequently quantifying the resulting changes in peak intensities.

**Figure 1.**
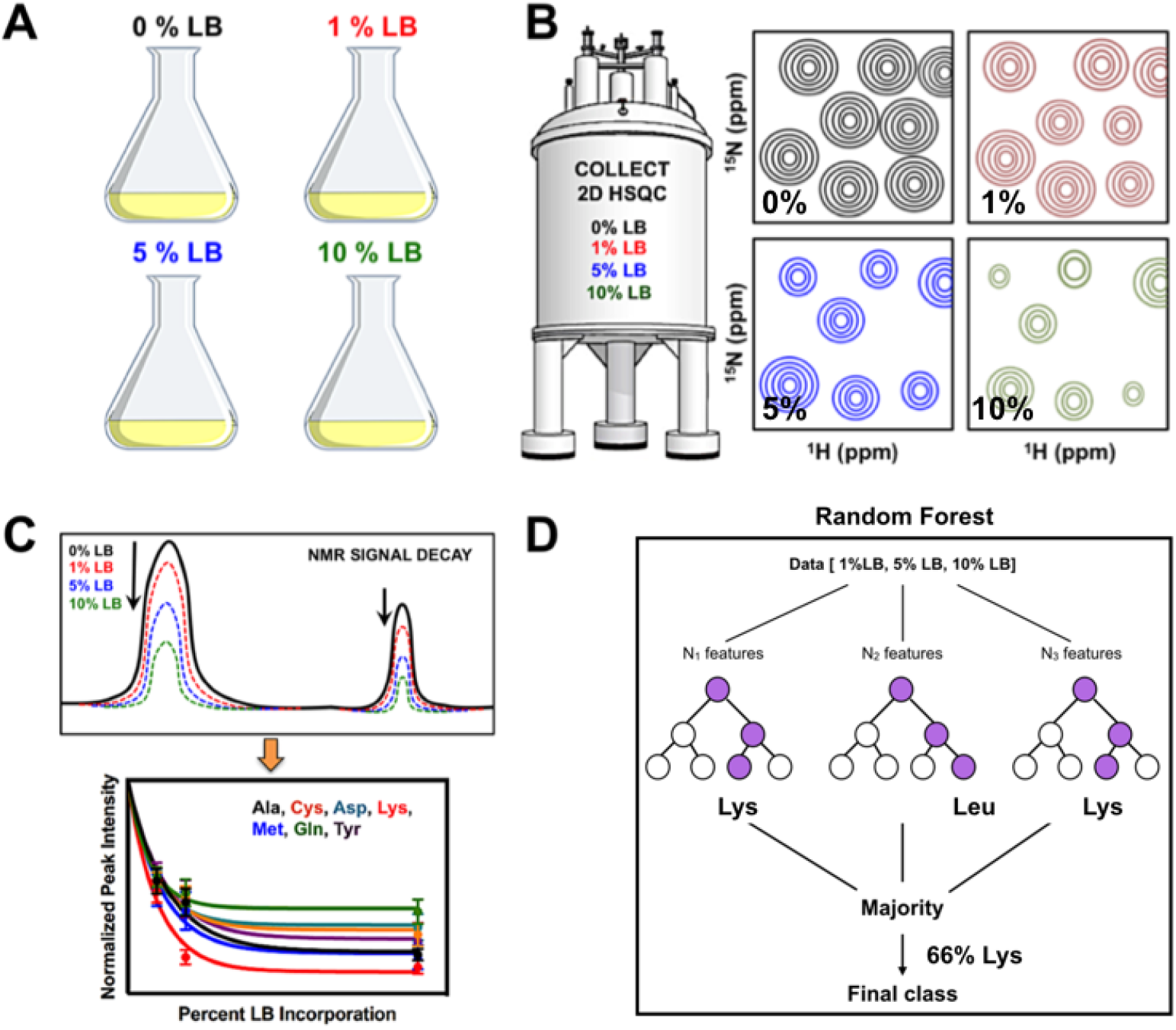
Graphical procedure for amino acid identification by LB doping. **(A)** Isotopically enriched (spin-1/2) M9 minimal medium cultures are doped with varying amounts of LB. **(B)** Cartoon of ^1^H-^15^N HSQC spectra of these samples show characteristic changes in resonance intensity. **(C)** NMR signal intensity decays are analyzed for each amino acid type. **(D)** A Random Forest algorithm is trained on the signal decays and is used for amino acid identification.

Our motivation for this work stemmed from the observation that for a number of different proteins the peak intensities in ^1^H-^15^N HSQC experiments varied throughout the spectrum, even in cases of rigid protein structures with isotropic rotational correlation times. In addition, the peak intensities were observed to vary between different samples of identical proteins. Upon further examination, we realized this was due to varying amounts of LB contamination that was present in the initial ‘starter’ cultures that were used to inoculate the large volume, ^15^N-labeled M9 growths. We hypothesized that depending on the amount of residual LB that was present in the M9 inoculation, ^14^N from the LB pool was being differentially incorporated into the amino acids via their biosynthetic pathways. Therefore, we attempted to exploit this phenomenon as a means to identify amino acids during the resonance assignment process. Taking advantage of the fact that glutamate (Glu) is a major nitrogen donor for bacterial amino acid biosynthesis (Fig 2) and the varied rates of aminotransferase activity and substrate inter-conversions throughout these biosynthetic pathways, the rate of ^14^N (and ^12^C) incorporation can be measured by the differential effect of added LB medium on signal attenuation in two-dimensional ^1^H-^15^N or ^1^H_3_-^13^C spectra.

**Figure 2.**
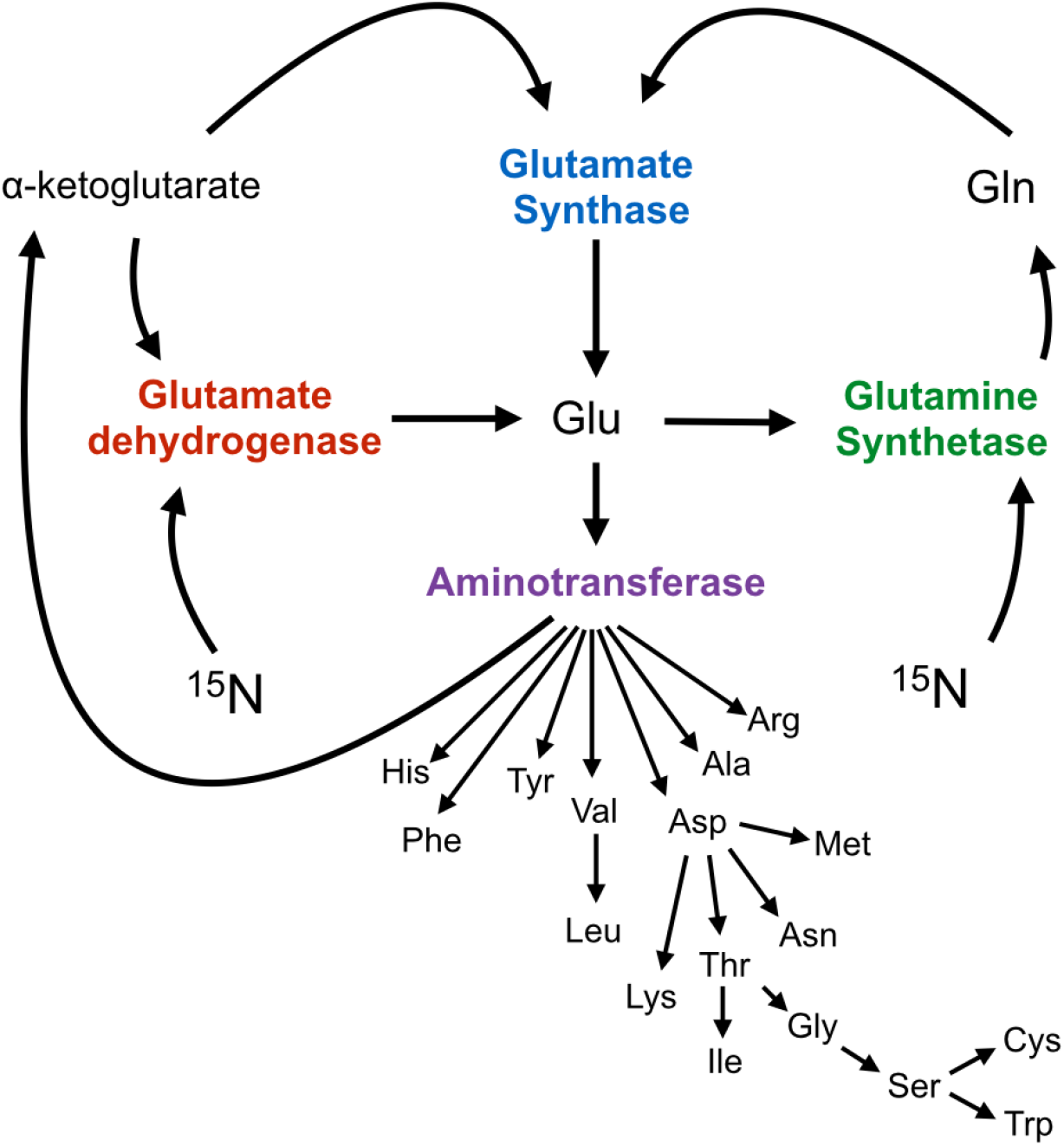
Schematic showing direction of nitrogen flow from glutamate to downstream amino acids. In the presence of excess ^15^N in the form of NH_4_Cl, glutamate dehydrogenase is responsible for the assimilation of ^15^N into glutamate through incorporation into α-ketoglutarate. Subsequently, ^15^N assimilation into all other amino acids occurs through aminotransferase activity involving glutamate and an appropriate α-ketoacid^45^.

Importantly, this method is capable of distinguishing several amino acid types as well as differentiating amino acids that commonly scramble with one another, which hampers other methods such as selective unlabeling^43^. This method can be particularly advantageous at the final stages of resonance assignment where the choices of amino acid assignments have been narrowed to a few remaining possibilities. Moreover, this method can also help resolve spectrally overlapped amide or methyl resonances, which can be useful in NMR spin-relaxation measurements, where peak overlap prevents accurate quantitation of signal decay. Using two large enzymes as test cases, imidazole glycerol phosphate synthase (IGPS, 51kDa) and protein tyrosine phosphatase 1B (PTP1B, 37kDa), we demonstrate that doping isotopically labeled M9 growth medium with small amounts of LB provides a new angle for tackling resonance assignment in large proteins and represents a useful technique to supplement traditional three-dimensional resonance assignment experiments.

## Materials and Methods

### Protein preparation

The PTP1B construct of amino acids 1-301 was transformed into BL21-CodonPlus (DE3)-RIL cells and expressed as previously described with the following modifications ^46^. Cells were grown in perdeuterated D_2_O (> 98% purity) M9 minimal medium supplemented with ^15^NH_4_Cl (1.0 g/L) and ^12^C glucose (2.0 g/L). Samples were prepared in which 0%, 1%, 5%, or 10% LB (v/v) in 100% D_2_O was doped into the 1L 100% D_2_O, ^15^N-labeled M9 culture at the start of the growth. The 500 ml of the cell culture was grown in 2L baffled bottom shaker flasks (180-200 rpm) at 37 ºC. Growth curves for 1% and 10% LB cultures are shown in Fig. S1. Cells were allowed to reach an OD_600_ of 0.8 – 1.0 before induction with 0.5 mM IPTG. Two post-induction test conditions were used for protein expression: 8 hours at 37 ºC and 20 hours at 20 ºC. After induction by IPTG, cells were pelleted by centrifugation and resuspended in a buffer containing 20 mM Bis Tris, 20 mM imidazole, 3 mM dithiothreitol (DTT), 10% (w/v) glycerol, 1 mM ethylenediaminetetraacetic acid (EDTA) at pH 6.5. Cells were lysed by ultrasonication, centrifuged, and the supernatant was applied to HiTrap Q HP and HiTrap SP HP columns (GE Healthcare) using a buffer of 20 mM Bis Tris, 20 mM imidazole, 3 mM DTT, 10% (w/v) glycerol, 1 mM EDTA, 0.5 M NaCl at pH 6.5 and eluted with a NaCl gradient. The PTP1B sample was concentrated to 0.18 - 0.3 mM (determined from A280, ε^280nm^ = 46410 M^−1^ cm^−1^) for NMR studies in a 50 mM HEPES buffer containing 150 mM NaCl, 0.5 mM TCEP, 7% D_2_O, and 0.03% NaN_3_ at pH 6.8.

The second test case we examined was IGPS, a non-covalently associated heterodimer comprised of two enzymes, HisH (23 kDa) and HisF (28 kDa). In the following LB tampering experiments, only HisF is isotopically labeled with spin-1/2 nuclei in the heterodimeric enzyme complex in which HisH is perdeuterated only. HisH and HisF are expressed separately and the *E. coli* containing these proteins are co-lysed to facilitate protein complex formation and purification ^47^. In the work described here, HisH is expressed in perdeuterated ^14^NH_4_Cl, ^12^C glucose, M9 minimal media, whereas HisF is expressed and isotopically labeled in BL21(DE3) RIL cells as previously described ^16^. The HisF protein was expressed in D_2_O (> 98% purity) ^15^N-labeled M9 minimal medium containing 0%, 1%, 5%, or 10% (v/v) of LB made up in 100% D_2_O, per liter. LB dopants were added at the initiation of bacterial growth, and cultures of HisF were grown at 37 °C with ^15^NH_4_Cl (Cambridge Isotope Labs, Tewksbury, MA) and ^12^C-glucose. Cells were allowed to reach an OD_600_ of 0.8 – 1.0 before induction with 1 mM IPTG. Cells were expressed under two post-induction conditions: 8 hrs at 37 °C, and 20 hrs at 20 ºC. The cells (containing isotopically labeled HisF and perdeuterated HisH) were harvested by centrifugation and resuspended in a buffer of 10 mM Tris, 10 mM CAPS, 300 mM NaCl and 1 mM β-mercaptoethanol at pH 7.5 and then co-lysed by ultrasonication. The cell lysate was clarified by centrifugation and the IGPS complex was purified by Ni-NTA affinity chromatography utilizing the C-terminal histidine tag on HisH. IGPS samples for NMR study were concentrated to 0.38 – 0.41 mM (determined from A280, ε^280nm^ = 29005 M^−1^ cm^−1^) in a buffer containing 10 mM HEPES, 10 mM KCl, 0.5 mM EDTA, and 1 mM DTT at pH 7.3.

A third enzyme, PHPT1 was expressed and purified to test the accuracy of amino acid prediction. The plasmid encoding the enzyme PHPT1 (14 kDa) was transformed into BL21(DE3) cells and grown in identical perdeuterated ^15^N-labeled M9 minimal medium cultures doped with 0, 1, or 10% (v/v) LB made up in 100% D_2_O. Cells were grown to an OD_600_ of 0.8 – 1.0 and following induction with 1 mM IPTG, grown an additional 16 hours at 25 °C and harvested by centrifugation. PHPT1 was purified by Ni-NTA affinity chromatography facilitated by a C-terminal histidine tag. NMR samples were concentrated to 0.17 – 0.29 mM in buffer containing 20 mM Bis-Tris, 50 mM NaCl, and 1 mM TCEP at pH 7.2.

### NMR Spectroscopy

NMR experiments were performed on a Varian 600 MHz spectrometer at 30 °C (IGPS), 25 °C (PHPT1), or 19 °C (PTP1B). ^1^H-^15^N TROSY spectra of IGPS, PTP1B, and PHPT1 were collected with the ^1^H transmitter and ^15^N offsets set to the water resonance and 120 ppm, respectively. HSQC experiments were collected with identical parameters including a 2 s relaxation recycle delay, 32 transients, 128 t_1_ increments, 4096 data points in the direct dimension, and spectral widths of 12000 Hz and 2800 Hz in the direct and indirect dimensions, respectively. The only exception was the 0% LB PTP1B sample in which the HSQC experiment was collected with 16 transients. NMR spectra were processed with NMRPipe^48^ with using apodization using a sine-bell window function to measure NMR resonance peak height. NMR spectra were analyzed in NMRFAM-Sparky ^49^. Changes in signal intensities between spectra were compared through the analysis of NMR resonance peak heights. The peak heights were determined in NMRFAM-Sparky through quadratic interpolation.

^15^N Longitudinal relaxation rates were determined using standard experiments^50^. Spectra were acquired with spectral widths of 1944 Hz and 12000k Hz in the t_1_ and t_2_ dimensions with 64 and 2k points, respectively. The proton carrier frequency was centered on the H_2_O signal and the ^15^N carrier set to 120 ppm. There are two sources of signal attenuation in the ^1^H-^15^N HSQC experiments described herein. The first is due to incorporation of ^14^N from the natural abundance nutrients in the intentionally added LB media. The second source of signal loss is due to enhanced relaxation of the amide nitrogen due to dipolar effects from the additional protons present in the LB that are subsequently incorporated into the proteins. In order for the proposed method to be useful the former source must be significantly larger and more varied than the additional ^1^H dipolar effects. The dominant relaxation mechanism for ^15^N amide is the directly bonded proton ^51^, however, in order to assess the effect of dipolar relaxation due to added protons originating from the LB media we performed two control experiments. The first experiment was a protein expression of PTP1B in ^15^N labeled perdeuterated M9 media with 10% by volume of 100% H_2_O with 90% D_2_O ^15^N-labeled M9 medium. The second experiment was a protein expression of IGPS in ^15^N labeled perdeuterated M9 media including 10% by volume of 100% ^1^H_2_O ^14^N-labeled M9 medium. ^1^H-^15^N TROSY spectra was collected for both samples and the peak heights of each spectra were analyzed and compared to the reference and LB doped spectra.

### Analysis of NMR resonances

For each assigned NMR resonance the peak heights were normalized to the protein concentration of the 0% LB sample using Eq (1). Here I_X_ represents the measured resonance signal intensities at x % LB addition, corrected for protein concentration relative to the 0% LB sample. I_N_ is the normalized intensity at N % LB, used in subsequent data analysis.

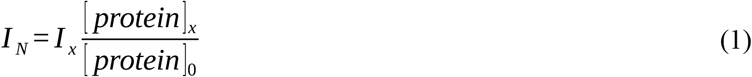

For comparison between different protein samples and expression times Z–scores of intensity ratios were calculated. For example, Z_10,1_ is calculated as in Eq (2):

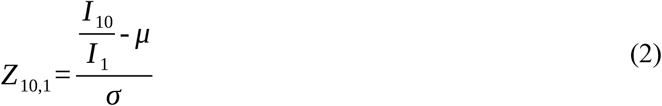

Here, μ is the mean of the normalized intensities for the entire data set and σ is the standard deviation. This analysis allows comparison of normalized intensity differences from the mean of all normalized intensities within a particular sample.

### Classification of NMR resonance assignments using Random Forests

In our analysis of the changes to NMR peak intensities from LB incorporation, we noticed differences in the magnitude of signal attenuation between different amino acids and different groups of amino acids. To aid in defining this relationship, we used the Random Forest classifier to group amino acids according to their NMR resonance decay profiles. Random forest classifiers can be described as an ensemble of decision trees (Figure 1D). A decision tree is essentially a flow chart of if-then-else clauses (for example “if signal is greater than 0.5, proceed to the left) which terminate in class identity. Decision trees are often fit through recursive partitioning of each variable, which determines the if-then-else clauses that yield minimal intra-class variance. Decision trees are often overfit to training data. To remedy this, random forests were invented, which are an ensemble of decision tress each fit from a random subsection of variables (columns) and samples (rows) ^52,53^. After training these *n* decision trees, unlabeled data is fed into each decision tree, each decision tree makes its guess at the class, and the prediction is determined to be the most popular class among all of the decision trees. This model is inherently multiclass and deals well with data variance, making it well-suited for identifying amino acid types ^54^. We used the scikit-learn implementation of this method in Python 3.7 environment ^55^. This implementation is ready-to-use, and only requires training data and parameters which dictate the way in which the model is fit (hyperparameters).

To construct our random forest model, we first selected non-overlapped NMR resonance peaks with S/N greater than 25 from the 8 hr-induction time PTP1B and IGPS datasets. We included 10 features or input variables as potential predictors for the random forest. These features and preprocessing included Z–scored natural logarithmic transformations of the 0%, 1%, 5%, 10% LB samples and Z–scored signal ratios of 1%/0%, 5%/0%, 10%/0%, 5%/1%, 10%/1%, 10%/5%. We then split the data (n = 269) into a 90/10 training (n = 242) and test set (n = 27). In order to determine the optimal hyperparameters (such as number of trees and depth of trees), we performed a random search with 5-fold cross validation on the training data. This technique involves splitting training data into 5 subsections, which are iteratively split 4 to 1 in order to yield 5 different evaluated models. It is useful to iteratively improve a model’s performance for two reasons: 1) it is the averaged performance between 5 different models, and hence does not reflect an over-fitted version of the model and 2) it allows for model tuning without contamination of the hold-out, test data. Cross validation was first performed in order to determine the number of decision trees to use in the model, which we determined to be 100. Next, this same process was used to determine the other following hyperparameters: tree depth, features per tree, the minimum number of samples required to split a node, whether to use the same peak in multiple trees (bootstrap), and function used to create tree splits. Parameters for each session of cross-validation were chosen by randomly sampling the following parameters 50 times: max depth (3 branches to none), maximum variables (1 to 10), minimum number of peaks to split a node (2 to 10), bootstrap (true or false), and tree split method (gini or entropy). This experiment yielded the following hyperparameters, which were subsequently used for each iteration of the model: max depth = 3, maximum variables = 4, minimum peaks to split node = 7, bootstrap = true, tree split method = entropy. Following optimization of the model, it was tested on hold-out test data using the following metrics: precision = TP/(TP+FP), model recall = TP/(TP+FN), f1 score = (1/2) * (precision * recall) / (precision + recall), and accuracy = TP/(TP+FP+FN), where TP = true positive; FP = false positive, FN = false negative. After training, we split the classifications of amino acids into 4 bins, (1) AILVHM, (2) DEQ, (3) NRSTFG, and (4) K, using the single-letter amino acid codes. Documentation of feature engineering, model optimization and framework can be found on github {https://github.com/evan-anderson/MeTA}

## Results

### Effects of LB Incorporation on ^1^H-^15^N HSQC Spectra of IGPS and PTP1B

Spectral overlap and relaxation effects in large proteins often make the resonance assignment process difficult. We used two model proteins, IGPS (HisF domain; 253 residues; MW = 28 kDa, (53 kDa in complex with HisH) and PTP1B (301 residues, MW = 37 kDa)), to demonstrate that doping small amounts of rich medium (LB) into protein expression protocols allows the metabolic variety of amino acid biosynthesis to become a tool to aid in their identification. The representative NMR spectra of IGPS and PTP1B shown in Fig. 3 illustrate that small amounts of natural abundance LB doped into the M9 growth alters the intensities of numerous resonances while leaving others nearly unperturbed.

**Figure 3.**
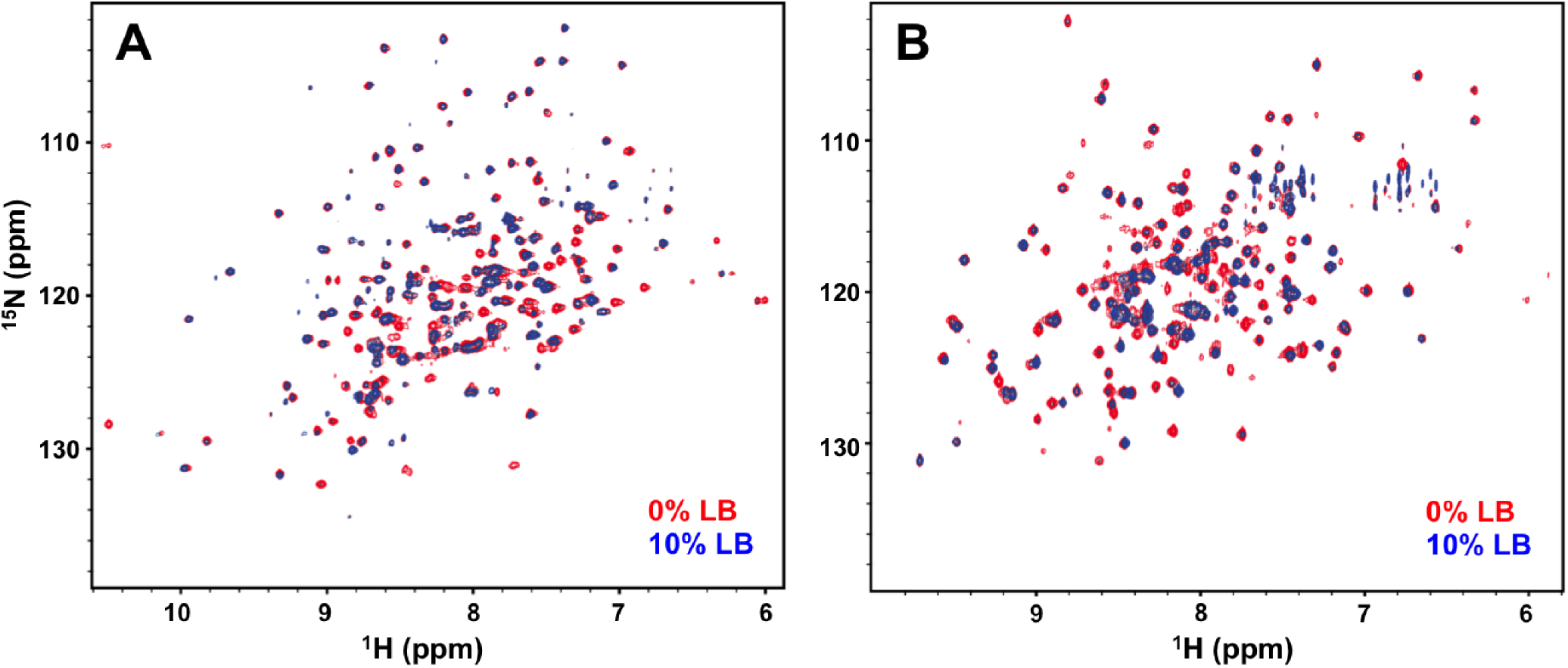
Representative ^1^H-^15^N TROSY spectra of IGPS **(A)** and PTP1B **(B)** grown and expressed in the absence (red) and presence of 10% LB growth medium (blue).

We analyzed the NMR signal intensity profiles for samples prepared with 1%, 5%, and 10% LB for PTP1B and IGPS. The normalized residual signal (I_1_, I_5_, I_10_) for PTP1B with the addition of LB at 1%, 5%, and 10% concentrations are shown in Fig. 4 A-C and profiles for IGPS are shown in Fig. S2. In PTP1B (IGPS), the addition of 1% LB resulted in an average loss of 45% (33%) signal compared to no addition of LB (I _1_/I_0_). There are no obvious differences between the amino acid types; all residues appear to show a close to uniform decrease in signal intensity at 1% doping. Upon incorporation of 5% and 10% LB, the profile of ^15^N/^14^N isotope scrambling between amino acid types become differentiable. The overall signal intensities (I_5_/I_0,_ I_10_/I_0_) of residues in PTP1B (IGPS) decreased on average by 75% (53%) with 5% LB and 85% (64%) with 10% LB incorporation. In the 5% LB NMR spectra of PTP1B (IGPS), the intensities of Lys resonances are noticeably reduced by 95% (90%). In the 10% LB spectra, ^15^N/^14^N isotopic scrambling becomes distinguishable between amino acids in different biosynthetic pathways. To highlight the relative differences, we show the I_10_/I_1_ for both PTP1B and IGPS in Fig 4D. The residual signal profile shows that along with Lys, Ala, branched chained amino acids (Ile, Leu, Val), Met, and His have the largest difference in signal between I_10_ and I_1_. Conversely, the smallest change in signal between 1% and 10% LB is observed for amino acids Asp, Glu, and Gln. It should be noted that there is only one Cys and one Trp in IGPS, and one assigned Met. The Met residue in IGPS was not considered in this sample, as it was overlapped with another peak.

**Figure 4.**
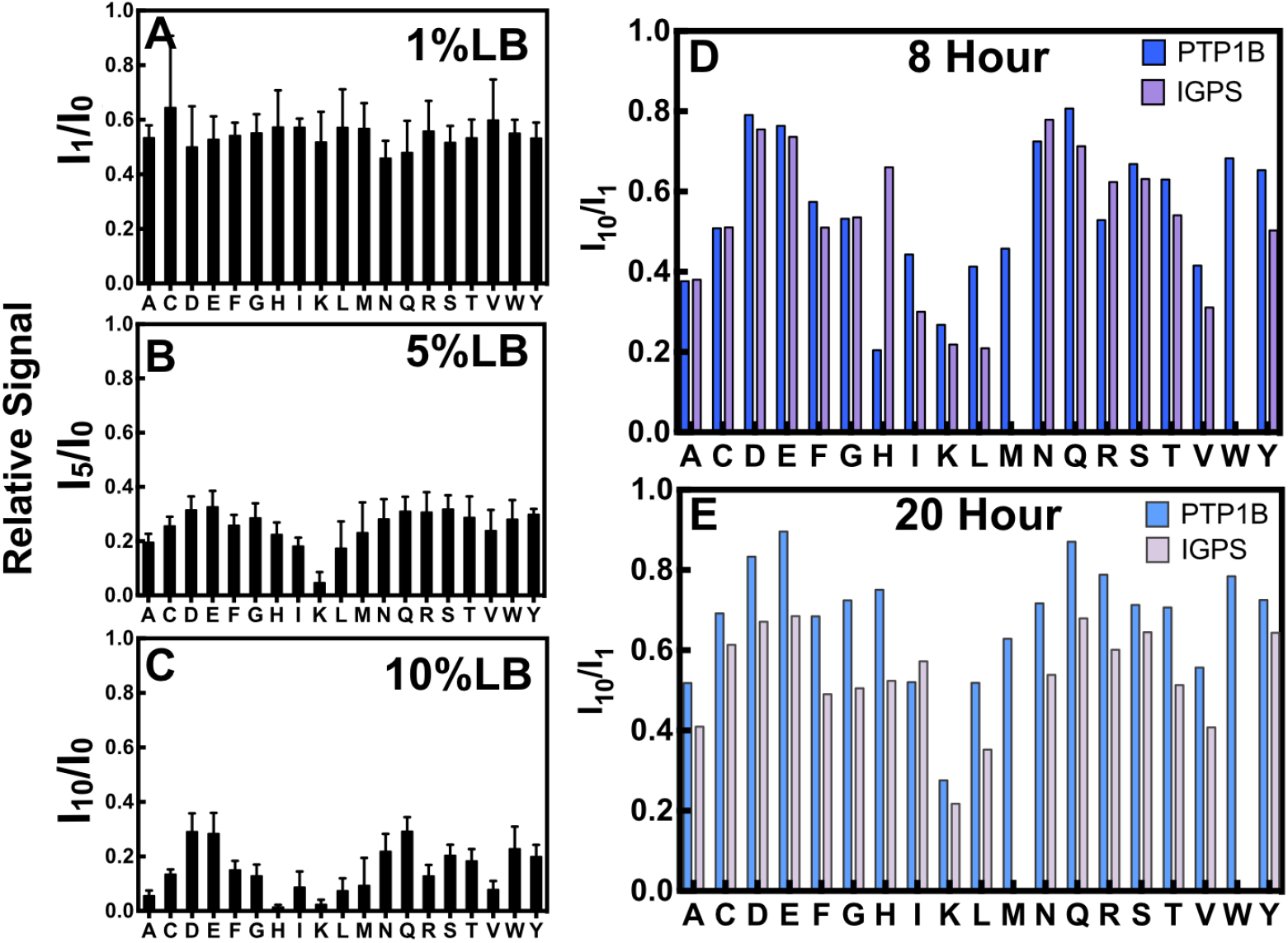
Average normalized peak intensity changes for amino acids in PTP1B in the presence of **(A)** 1% LB, **(B)** 5% LB, and **(C)** 10% LB for an 8 hour expression determined from a ^1^H^15^N TROSY HSQC. Plots are analyzed as an average of all resonances of a specific amino acid type. The ratio of residual signal from the 10% LB spectra over 1% LB spectra for **(D)** 8 hour expression and **(E)** 20 hour expression shows changes in the amino acid signal for PTP1B and IGPS.

In addition to the 8 hour growth, we show the 1% LB and 10% LB signal intensity differences for PTP1B and IGPS in a 20-hour expression experiment in Fig. 4E. The results revealed a similar to trend to that observed in the 8-hour experiment, where, Lys, Ala, Leu, Ile, and Val exhibit large signal intensity decreases, while the Asp, Glu, and Gln residues show higher signal retention at I_1_/I_0_ and I_10_/I_0_. These results indicate that the degree of ^15^N/^14^N isotopic scrambling does not depend on expression times, but rather the concentration of LB present in the growth media.

One source of this signal attenuation, which could occur through incorporation of ^1^H into the protein from the natural abundance LB media is ^1^H dipolar relaxation. Control experiments (Fig. S3 and S4) with 10% H_2_O show that the dipolar-based contribution to signal attenuation for PTP1B can be estimated to be 26% ± 12%. Additionally, an experiment with IGPS expressed with 10% (H_2_O, ^14^N) showed reduced signal intensity by 36% ± 12% – this accounts for the 10% decrease in signal due to incorporation of ^14^N and 26% signal attenuation due to ^1^H dipolar relaxation effects. *T*_*1*_ measurements for 10% (H_2_O, ^14^N) and 10% LB IGPS and T_2_ estimations of control experiments and 10% LB doped experiments are shown Fig. S5 and S6. Both T_1_ and T_2_ line width analysis reveals that the 10% H_2_O control is a good approximation of the contribution of ^1^H dipolar relaxation. It is worth noting that in the 10% LB experiments, we observe signal reduction greater than 36% for all amino acid types, which indicates that we are not observing uniform ^14^N/^15^N scrambling. Therefore the variability in signal observed across the amino acids with LB dopant (Fig. 4) are due to distinctions in the amino acid metabolic pathways.

Additionally, we wanted to examine the possibility that the signal attenuation may be dependent on the concentration of amino acid levels present in LB. We predicted 5% and 10% LB relative peak intensity values by extrapolating from 1% IGPS data, and compared it to the real 5% and 10% LB IGPS data set (Fig. S7). We found that only the peak intensities of Lys residues could be predicted, indicating that signal attenuation for Lys residues is concentration dependent. All other amino acids types had relative signal intensities greater than the predicted value. We believe this can be attributed to the balance between ^15^N donation through the glutamate / transaminase activity in amino acid biosynthetic pathways and the different mechanisms of amino acid interconversion between ^14^N metabolites that are introduced into the bacterial growths in the D_2_O LB medium. This hypothesis is reinforced by observing the effect of LB doping on ^13^CH_3_ methyl labeling of Ile, Val, and Leu residues in TROSY based ^1^H-^13^CH_3_ HMQC spectra shown in Fig. S8 ^56^. IGPS growths were supplemented with ^13^CH_3_ α-ketoisovaleric acid and α-keto butyric acid to label ^13^C methyl of Leu, Val, and Ile residues respectively. We observed that the signal intensities of Leu residues are strongly affected at 10% LB and disappear from the ^13^C methyl spectra, while both Ile and Val retain partial signal intensity. Both Leu and Val residues are synthesized from the same precursor: α-ketoisovaleric acid. The asymmetrical change in signal intensity between Leu and Val is evidence of selective product inhibition for the Leu biosynthetic pathway and not the Val synthetic pathway.

### Z-score to Compare Data Sets

In these experiments, Lys residues are easily identified with 5% LB dopant, as they are the only peaks that lose over 90% signal intensity. The identification of the other amino acid types presents as a more complicated problem, as they have varied degrees of peak attenuation between 1–10% LB. Additionally, notable variance is observed when evaluating the absolute peak intensity changes between the IGPS and PTP1B 8hr datasets. A number of extrinsic factors could be the cause of signal variation between PTP1B and IGPS. For example, variation in the pre-induction growth time or rates of protein production can attribute to these differences. Although the absolute values between the two test proteins are different, we observed a pattern in the relative signal changes between amino acid types. This pattern becomes apparent with the calculated z-scores of the peak intensity ratios for each amino acid type and can be extended to data collected at 20 hrs. The peak intensities ratios are shown to have a normal distribution (shown in Fig. S9), enabling z-score analysis. We show the boxplots of combined 8 hr and 20 hr data for Z(I_10_/I_0_) and Z(I_10_/I_1_) compared between PTP1B and IGPS in Fig. S10. Both proteins follow the same trend in signal attenuation, with more distinction from Z(I_10_/I_1_). Figure 5A and Table 1 shows the average z-score for Z(I_10_/I_1_) for each amino acid 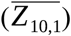, and its standard deviation (*σ* _*Z*_) for combination of the 8 hr and 20 hr datasets of PTP1B and IGPS.

**Table 1.** Z(I_10_/I_1_) values calculated for 8 and 20 hr PTP1B and IGPS data set.

| | $\overline{Z}_{10,1}$ | $\sigma_{\overline{Z}}$ | $\overline{Z}_{10,1} \pm 2\sigma_{\overline{Z}}$ |
| --- | --- | --- | --- |
| **Ala** | -0.78 | 0.25 | -0.27 to -1.28 |
| **Cys** | 0.14 | 0.27 | 0.69 to -0.40 |
| **Asp** | 0.98 | 0.53 | 2.03 to -0.07 |
| **Glu** | 1.12 | 0.73 | 2.58 to -0.34 |
| **Phe** | -0.07 | 0.31 | 0.54 to -0.68 |
| **Gly** | -0.07 | 0.34 | 0.61 to -0.75 |
| **His** | -0.30 | 0.98 | 1.67 to -2.26 |
| **Ile** | -0.65 | 0.40 | 0.15 to -1.45 |
| **Lys** | -1.39 | 0.41 | -0.57 to -2.21 |
| **Leu** | -1.06 | 0.50 | -0.07 to -2.05 |
| **Met** | -0.57 | 0.58 | 0.58 to -1.73 |
| **Asn** | 0.41 | 0.62 | 1.65 to -0.82 |
| **Gln** | 1.00 | 0.48 | 1.95 to 0.05 |
| **Arg** | 0.18 | 0.78 | 1.73 to -1.37 |
| **Ser** | 0.46 | 0.56 | 1.58 to -0.65 |
| **Thr** | 0.11 | 0.33 | 0.76 to -0.55 |
| **Val** | -0.76 | 0.46 | 0.17 to -1.68 |
| **Trp** | 0.54 | 0.47 | 1.48 to -0.41 |
| **Tyr** | 0.32 | 0.33 | 0.99 to -0.35 |

**Figure 5.**
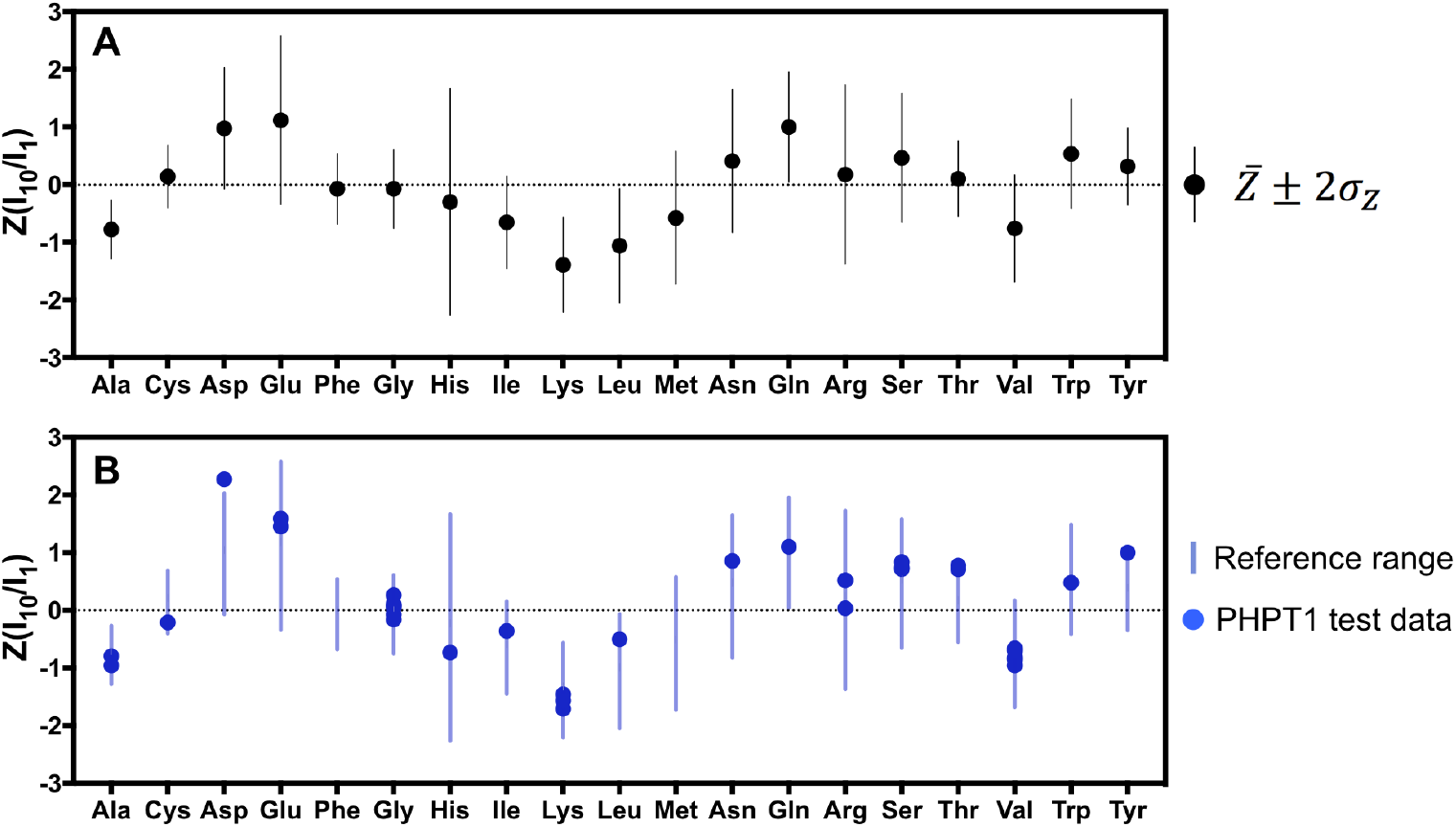
Z–score analysis of the signal intensity differences between 1% and 10% LB experiments. **A)** The 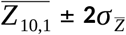 values for Z(I_10_/I_1_) for each amino acid type obtained from 8 hr and 20 hr expression experiments of PTP1B and IGPS. **B)** Z(I_10_/I_1_) values for the PHPT1 test data (34 peaks) expressed at 25ºC for 16 hr compared to the reference range 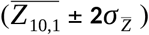 determined in **A)**. The vertical bars represent 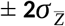 values from the 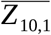.

The z-score of the data show a trend between the amino acid types due to the attenuation of the NMR resonances by the addition of LB. For example, there is a clear separation between amino acids types that are insensitive to LB addition (Glu, Asp, and Gln), and amino acid types that have high signal attenuation (Ala, Leu, and Lys) (Fig. 5A). Additionally, we observe His, Arg, Asn, and Glu have larger variability between different growth conditions as indicated by the larger vertical bars in Figure 5. In contrast, the range of ^14^N/^15^N incorporation of other residues such as Ala, Cys, and Phe is much more consistent across samples.

To demonstrate that the Z(I_10_/I_1_) calculated here is robust and can be applied to other protein systems and different growth conditions, we collected NMR spectra of 1% LB and 10% LB-doped ^15^N-labeled PHPT1 (14 kDa) grown at 25 ºC for 16 hours. A total of 36 non-overlapping peaks were selected as the test sample. The Z(I_10_/I_1_) of PHPT1 peaks were calculated and compared to the observed experimental ranges determined from PTP1B and IGPS data (compare patterns in Fig. 5A and B). We found that 34 out of 36 peaks (94%) in PHPT1 had Z(I_10_/I_1_) values within the reference range of the corresponding amino acid type determined from the PTP1B/IGPS data set. The table of the calculated Z(I_10_/I_1_) values PHPT1 test data set is found in Table S1. The observed outlier was residue D58, which had an observed Z(I_10_/I_1_) of 2.30, slightly out of the reference range (µ ± 2σ) for Asp (-0.07 to 2.03). The reproducibility of these Z(I_10_/I_1_) values at different temperatures (37 ºC, 25 ºC, and 20 ºC) and expression times (8 hr, 16 hr, and 20hr) demonstrates that that the measured signal intensity differences between the amino acid types are due to the inherent differences in amino acid biosynthesis rates.

### Random Forest for Amino Acid Classification

Although we have found a way to distinguish amino acid types with only the analysis of 1% LB and 10% LB ^15^N labeled spectra, the variance in the I_10_/I_1_ data hampers the discrimination of NMR resonance assignments with Z(I_10_/I_1_) values near the mean (-0.7 to 0.7) in PHPTI (Fig. S11). In an effort to create a reliable tool for rapid categorization of amino acid types and enhance the ability to differentiate amino acids with degenerate Z(I_10_/I_1_) values, we decided to use a supervised machine learning algorithm – random forests ^52^, which is a robust classification method used in many areas of biology ^52,57^. We sought to improve the differentiation of amino acid types by training the model on the 8 hr PTP1B and IGPS NMR resonance datasets (n = 282), integrating observed changes from 0%, 1%, 5%, and 10% LB doping. We also included additional features to describe the signal decay relationship between datasets: 1%/0%, 5%/0%, 10%/0%, 5%/1%, 10%/1%, and 10%/5%. The Z–scores were calculated for all the features to enable comparison between protein samples. To ensure model accuracy, we only included NMR resonance peaks that had good signal to noise and were non-overlapped. Additionally, we removed amino acids where the total representation was < 5: Trp, Tyr, Cys. The final dataset (n = 269), was then randomly split into training (90%) and test (10%) sets, and we then used training data under 5-fold cross-validation to evaluate base performance of the scikit-learn random forest classifier ^55^. Subsequently, we altered data preprocessing and model hyperparameters in order to yield the best model score through cross-validation. Based on the observed Z(I_10_/I_1_) ranges, the prediction buckets were split into four amino acid categories: (1) Lys (K); (2) Ala, Ile, Val, Leu, Met, His (AILVMH); (3) Asn, Arg, Gly, Phe, Ser, Thr (NRGFST), and (4) Asp, Glu, Gln (DEQ). The predictions of the training data using this model were applied to the test data set and yielded 25 true positives out of 27 total, or a model accuracy of 92%. A matrix summarizing the ability of the model to correctly label test data (confusion matrix) is shown in Fig. 6A. Additionally, the precision and recall calculated for each category are summarized in Fig. 6B. High precision and recall across all groups in the test dataset means that A) the model is not overfit to training data and B) the model has not overfit to a specific group. Two additional models were built where one model was trained on PTP1B (n = 147) and tested on IGPS (n = 122), and the other was trained on 80% (n = 215) of the combined datasets and tested on 20% (n = 54) of the hold-out data. We found the model that was trained on PTP1B and tested on IGPS resulted in 74% accurate predictions, and for the 80%/20% training/test model, the accuracy was 80%. The confusion matrices of both of these models are shown in Fig. S12. The results indicate that the accuracy of the MeTA prediction tool is data-limited: as more is used to train the model, the predictive power of the model on hold-out data increases.

**Figure 6.**
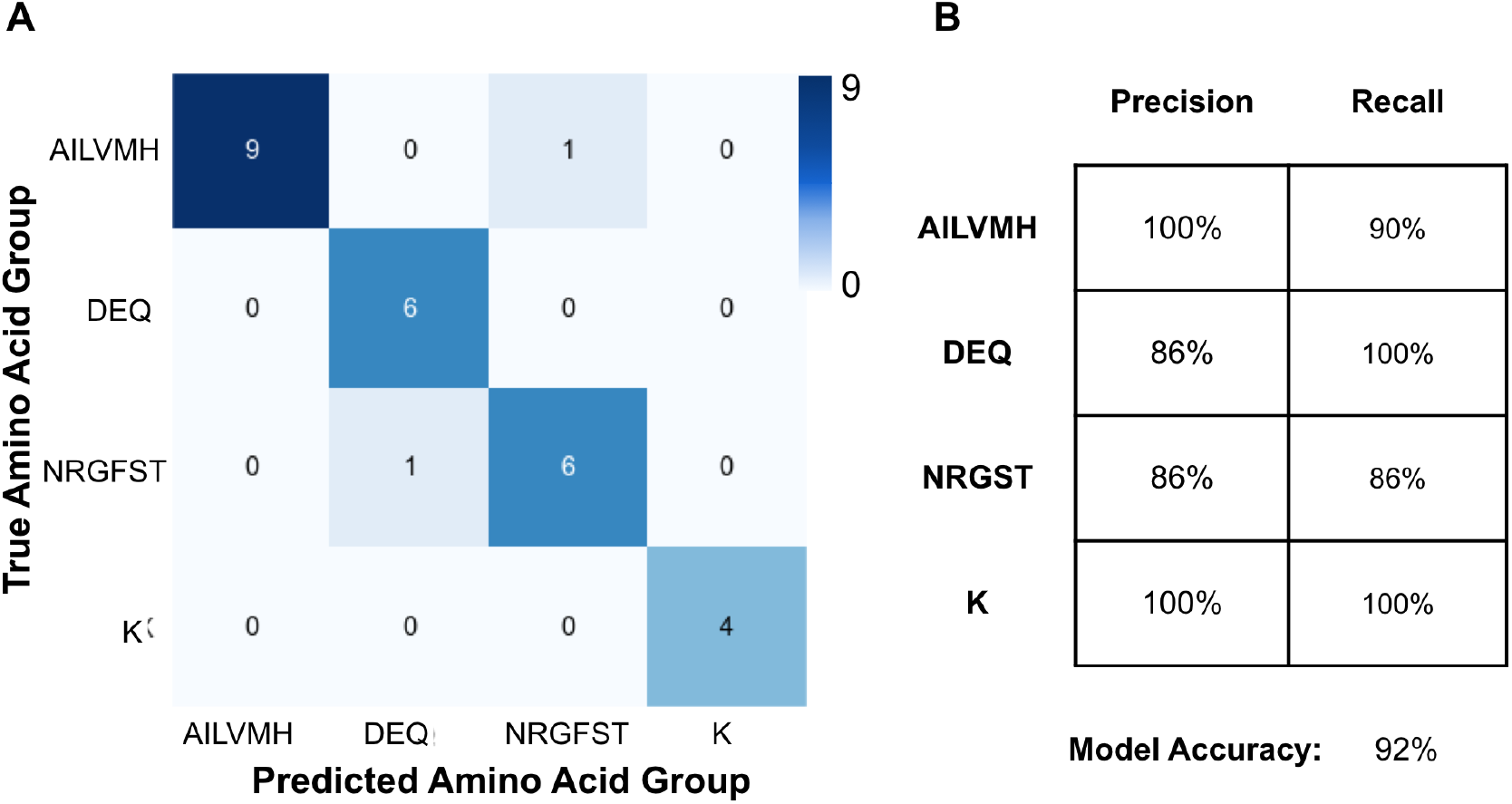
Confusion matrix summarizing the result of a metabolic tampering for amino acid (MeTA) prediction tool generated with the scikit-learn random forest algorithm. The model was trained on 8 hr PTP1B and IGPS datasets. **A)** Test data set showing prediction results classifying NMR resonances (n = 27) into AILVHM, DEQ, NRGFST, or K categories. **B)** Table summarizing the precision and recall values of the classification groups.

## Discussion and Conclusions

Identification of amino acids in NMR spectra of large proteins is complicated by high peak density resulting in unresolved resonances for many residues in three-dimensional experiments. It is therefore critical to devise creative ways to disentangle and assign these spectra for molecular insight, and advances in selective isotope incorporation and NMR pulse sequences have assisted the spectral assignment process. However, there remains room for improving on the limitations of many currently available assignment methods, many of which require preparation of numerous expensive samples or long experiment times. For example, resonance assignment by site-directed mutagenesis techniques is labor and sample intensive and risk alterations to the overall structure of the target protein. Specialized experiments detecting specific amino acid types in large proteins ^58–61^ can suffer from diminished sensitivity and are often ineffective for deuterated samples that do not retain side chain protons^43^. Isotope labeling schemes such as MILVAT (**<u>M</u>**et-**<u>I</u>**le-**<u>L</u>**eu-**<u>V</u>**al-**<u>A</u>**la-**<u>T</u>**hr) ^4^are tremendously useful for identifying methyl-containing side chains, but are not useful for more traditional ^1^H-^15^N HSQC spectral assignment and also require higher dimensionality (*i*.*e*. 4D) experiments that can strongly attenuate signal-to-noise. Further, the nature of these experiments preclude assignment of methyl groups that are not part of a methyl-methyl network, detectable by NOEs, from being identified ^4^. Other clever methods such as fractional ^13^C labeling uses various ^13^C carbon sources (acetate, [2-^13^C]- or [1,3-^13^C]-glycerol, [3-^13^C]-pyruvate, or [1-^13^C]-glucose) and cellular metabolism to selectively incorporate ^13^C labeling patterns for side chain amino acid identification^62–67^.

These approaches have been useful for NMR relaxation studies. More recent efforts have demonstrated the versatility of leveraging amino acid metabolism for resonance assignment applications ^44,68^. Meier and coworkers have used a similar approach of examining interconnected metabolic pathways to devise more efficient site-specific labels for solid-state NMR (ssNMR) applications. Addition of various combinations of labeled (^13^C, ^15^N) and unlabeled (^12^C, ^14^N) amino acids to bacterial cultures were shown to decrease spectral overlap in studies of a large membrane protein and facilitate resonance assignment by a series of dipolar assisted rotational resonance (DARR) experiments that maintain better sensitivity than traditional ssNMR N-C_α_-CO correlation measurements ^69^ A related study by Arthanari, Wagner and coworkers used 2-^13^C-pyruvate and 3-^13^C-pyruvate as bacterial carbon sources to selectively label ^13^C_α_ and/or ^13^C_β_ positions of amino acids ^68^.The critical role of pyruvate in the direct or indirect biosynthesis of nearly all amino acids was shown to generate unique signatures that enhanced the resolution of the standard HNCA such that it can serve as the workhorse triple-resonance experiment for assignment. Another useful approach, selective unlabeling, has been employed to reduce spectral complexity ^70,71^and assign resonances by incorporating a subset of amino acids in unlabeled form (via ^12^C_6_H_12_O_6_ and ^14^NH_4_Cl) into bacterial cultures, resulting in the disappearance of those amino acids from two-dimensional spectra and in their triple-resonance correlations. However, applications of this method can require many samples, depending on the amino acids of interest, and the utility of selective unlabeling is hampered by natural misincorporation of and cross-metabolism between related amino acids, precluding unambiguous identification of several residue types.^43^ Our approach is yet another way to effectively hijack bacterial metabolism to generate unique signatures for amino acids that bolster the utility of two-dimensional NMR spectra. We show that by accounting for metabolic regulatory mechanisms such as product inhibition, and differing anabolic and catabolic rates of amino acid synthesis, commonly scrambled subsets can be identified with as few as three 2D ^1^H-^15^N HSQC experiments. Specifically, we observed that amino acid resonances have distinct rate of signal attenuation following short (*i*.*e*. 8 hr, 37 ºC) or long (*i*.*e*. 20 hr, 20 ºC) bacterial growths for different LB dopant concentrations. By only collecting two HSQC spectra with 1% and 10% LB dopant, we show that it is possible to separate amino acid types with low signal attenuation such as Asp, Glu, and Gln from amino acid types that have high signal attenuation under known LB concentrations, such as Ala, Leu, Lys, Ile and Val. We have presented a table of Z(I_10_/I_1_) values corresponding the amino acid type (Table 1). The z-score analysis of peak intensity facilitates the comparison of changes in peak intensities for each amino acid type between protein systems in a protein size and concentration independent manner. The relative peak intensity differences are described in units of standard deviation relative to the mean I_10_/I_1_ value for the protein. In addition, we have tested these Z(I_10_/I_1_) ranges against a third protein PHPT1 which was grown at 25 ºC for 16 hr. In this test dataset, 97% of the Z(I_10_/I_1_) values fell within the reference values in Table 1. It is evident that this phenomenon holds true regardless of expression times and temperatures, and the Z(I_10_/I_1_) values presented in this study are robust and can be used for amino acid assignment or validation purposes. Particularly, the Z(I_10_/I_1_) metric will be useful at the end of the protein backbone assignment process, where there will be a few unassigned resonances, that can not be distinguished by other triple resonance experiments.

Additionally, we demonstrate through application of supervised machine learning, that it is possible to further classify NMR resonances into four amino acid categories: AILVHM, DEQ, NRGFST and K categories. The random forest algorithm was selected because it is low in computational cost, while robust to outliers and data variance, making it an ideal method for data classification. The model includes features obtained from 0%, 1%, 5%, and 10% 8 hr IGPS and PTP1B 2D ^1^H-^15^N HSQC data. The accuracy of the current model is limited by the size of the training data (n = 269). Future efforts in data collection for LB tampering experiments will allow for improvements in model accuracy and robustness. We believe that the method developed in this work using metabolic tampering for amino acid (MeTA) prediction will improve the timeline of NMR assignment due to the low cost in sample preparation and experiment time. In particular, one of the advantages of MeTA prediction is the ability to differentiate amino acid residues with degenerate Cβ shifts (Fig. S13) ^72^. For example the overlap of Cβ shifts for Asn and Asp (Fig S5) can hinder assignment from HNCACB experiments but these residues are easily differentiated using the method described here. This is also especially true for amino acids with Cβ shifts degenerate with those of Lys. Additionally, as far as we know, this is the only method that can selectively identify Asp, Glu, and Gln backbone amide chemical shifts, residues that often contribute to enzyme active sites.

The method reported herein presents a new avenue to rapidly identify amino acid types in the NMR spectra of large proteins (>25kDa) and supplement conventional 3D experiments most often used in NMR resonance assignment. Analyses of simple two-dimensional experimental data provide a fingerprint of each amino acid found in ^1^H-^15^N HSQC and ^13^CH_3_ HMQC spectra. The amino acid biosynthetic pathway can be leveraged to aid in the identification of residue types in NMR spectra by doping small amounts of LB into D _2_O M9 minimal medium bacterial cultures. The anabolic and catabolic pathways of amino acids stemming from the TCA cycle and glycolytic pathways all have distinct rates of interconversion, which is evident in data collected over short (8 hr) and long (20 hr) bacterial growth periods. Despite the varied expression times and temperatures of our model proteins, the observed pattern of NMR signal attenuation is distinguishable for each amino acid type, highlighting this approach as applicable and customizable to any bacterial growth system and providing unique flexibility to study large proteins with varied expression conditions.

This method is highly synergistic to protein NMR chemical shift prediction tools,^73–75^where, assignment of amino acid type using our method coupled to chemical shift predictions based on structural coordinates could guide sequential backbone assignment efforts in the absence of three-dimensional assignment data ^76^. Additionally, application of LB tampering can be used for relaxation studies. We observed that 30% of NMR resonance peaks have sufficient S/N quality for relaxation dispersion studies in the 10% LB PTP1B dataset (Fig. S14). Crowded overlapped regions can become resolved with LB doping (Fig. 7), therefore making it an appealing strategy to study dynamics in large proteins. As demonstrated in Figure 7, accurate quantification of peak intensities for R47 during a relaxation series would be complicated by overlap with K237. However, the rapid decay of signal for Lysine residues due to LB incorporation can be exploited to resolve R47 as shown in Figure 7 and thus make its peak quantitation straightforward.

**Figure 7.**
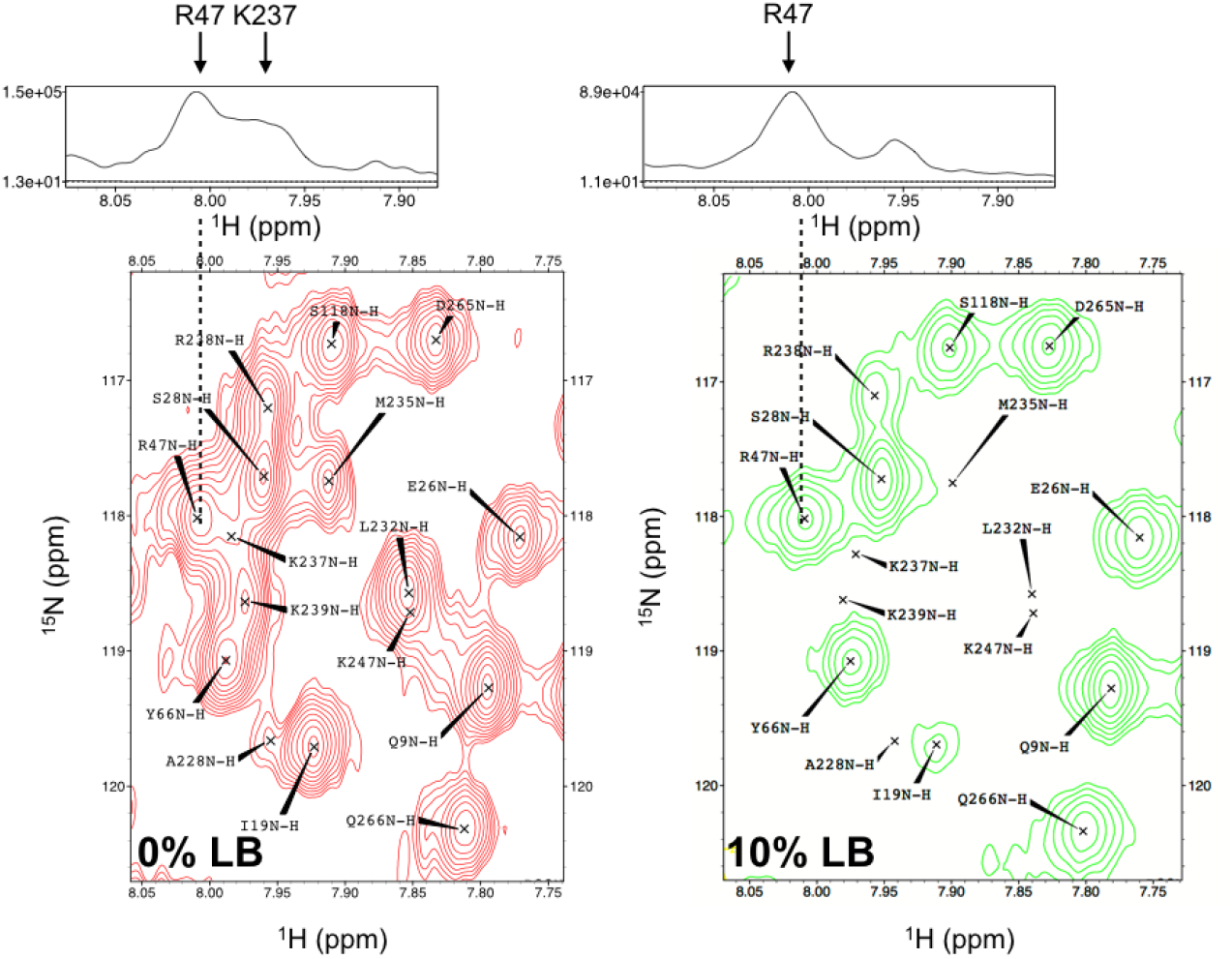
A comparison of a spectrally crowded region in 0% (red) and 10% (green) LB. (top) one-dimensional ^1^H slice is shown for R47 in the 0% and 10% LB spectra. The disappearance of K237 peak in the 10% LB spectrum enables the resolution of the resonance for R47. The S/N for R47 in 10% LB spectrum is 47.

This work draws attention to the sensitivity of metabolic scrambling and its potential to affect NMR spectral quality and signal intensity with minimal LB contamination, while also creating an avenue to supplement traditional resonance assignment with simple two-dimensional NMR data for proteins. The python script developed for MeTA prediction will be available in the Jupityr Notebook on github (https://github.com/evan-anderson/MeTA), along with the reference 8 hr PTP1B and IGPS dataset. A scheme of user input data and work– flow is shown in Fig. S15. The user will have the option to evaluate experimental data against the model trained on the datasets described within, as well as have the option to include their own dataset in the training process.

## Acknowledgments

This work was supported by National Institutes of Health grants GM106121 and GM112781, and NSF MCB1615415 to JPL

