## Supplemental Figures for "Facilitating NMR Resonance Assignment with Metabolic Tampering"

**Table of Contents**

**Figure S1:** O.D. growth curve of *E.coli* in different combinations of LB and M9 media

**Figure S2:** Summary of normalized peak intensity changes for amino acids in IGPS

**Figure S3:** Control experiment for the effect of $^{1}$H-induced relaxation on peak intensities for IGPS

**Figure S4:** Control experiment for the effect of $^{1}$H-induced relaxation on peak intensities for PTP1B

**Figure S5:** $T_1$ measurements for 10% ($H_2O$, $^{14}$N) and 10% LB IGPS

**Figure S6:** $T_2$ estimations from line width analysis

**Figure S7:** Extrapolation of peak intensity changes from 1% IGPS data

**Figure S8:** Effects of LB doping on $^{13}CH_3$ HMQC NMR spectra of Ile, Leu, and Val methyl groups in IGPS

**Figure S9:** Empirical distribution function of signal intensity ratios in IGPS and PTP1B

**Figure S10:** Boxplots of $Z(I_{10}/I_0)$ and $Z(I_{10}/I_1)$ values for PTP1B and IGPS

**Figure S11.** PHPT1 test data amino acid hits based on $Z(I_{10}/I_1)$ values

**Figure S12:** Confusion matrices of additional random forest models for amino acid classification

**Figure S13:** Comparison of Z-scores for residues with overlapping $^{13}$Cβ chemical shifts

**Figure S14:** The signal to noise of NMR resonance peaks observed in 10% LB $^{15}$N HSQC

**Figure S15:** Schematic of the user–end process on using random forest MeTA prediction tool

**Table S1:** $Z(I_{10}/I_1)$ values of PHPT1 test data (n=34) . Protein was expressed at 25ºC for 16 hr.

**SI Methods**

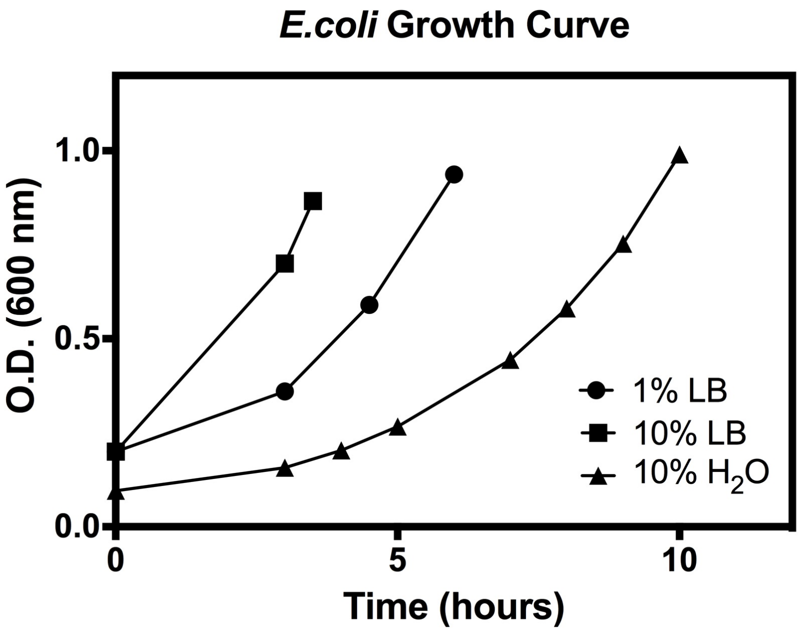


**Figure S1.** O.D. growth curve of *E.coli* BL21-CodonPlus (DE3)-RIL cells in $D_2O$ M9 minimal media with 1% LB, 10% LB, and 10% $H_2O$.

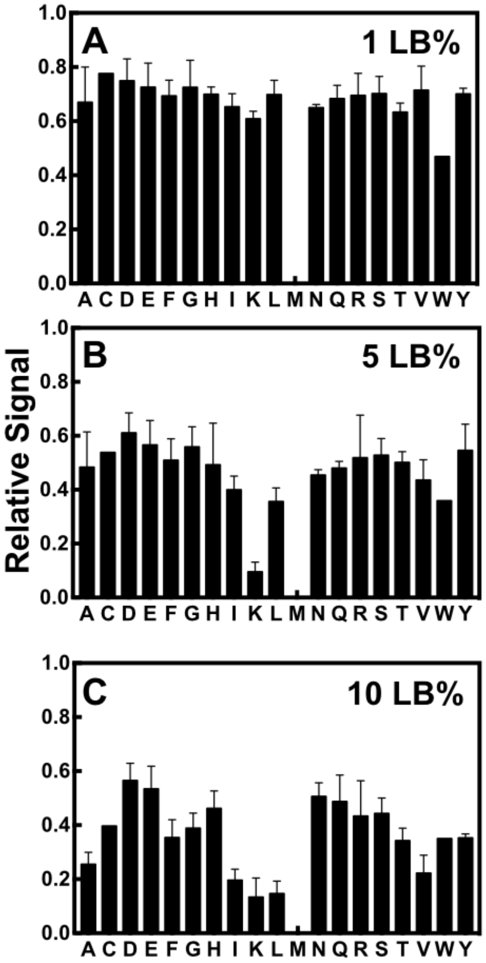


**Figure S2.** Summary of normalized peak intensity changes for amino acids in the HisF subunit of IGPS determined from $^{1}H$-$^{15}N$ TROSY experiments in which 1% **(A)**, 5% **(B)**, or 10% **(C)** LB medium was incorporated into the bacterial culture.

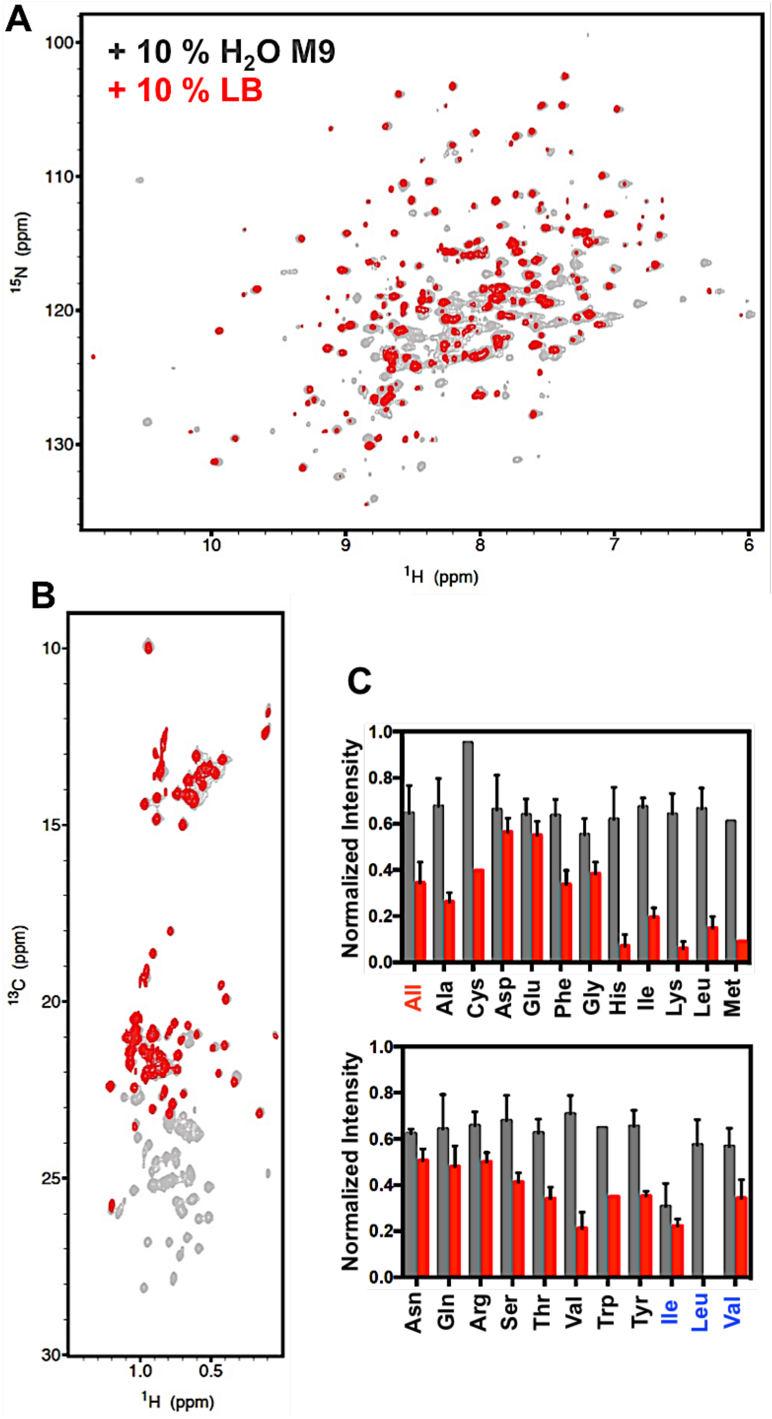


**Figure S3.** Control experiment (expressed for 8hr at 37ºC) assessing the contribution of $^{1}$H-induced dipolar relaxation to the attenuation of NMR signal intensities. **(A)** $^{1}$H-$^{15}$N TROSY spectral overlays comparing identical samples of IGPS where 10% by volume of 100% ($^{1}H_2O$, $^{14}$N) -M9 minimal mediums (gray) or 10% LB medium (red) was incorporated into the bacterial protein expression culture. **(B)** $^{1}$H-$^{13}CH_3$-ILV HMQC spectral overlays comparing the same experimental conditions described in **(A)**. **(C)** Summary of residual peak intensities for every non-proline amino acid in the presence of 10% $H_2O$ M9 (gray) and 10% LB (red). Values are also reported for the entire data set (red text) and for Ile, Leu, and Val residues in $^1H$-$^{13}CH_3$ HMQC experiments shown in **(B)** (blue text)

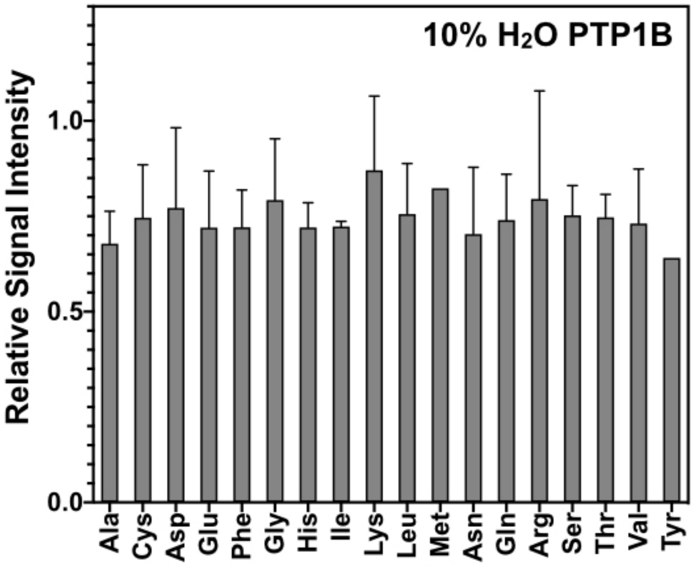


**Figure S4** – PTP1B control experiment (expressed for 8hr at 37ºC) assessing the contribution of $^1H$-induced dipolar relaxation to the attenuation of NMR signal intensities. Summary of relative peak intensity in the presence of 90%$D_2O$/10% $H_2O$ M9 compared to $^{15}N$ labeled PTP1B grown in 100% $D_2O$ M9.

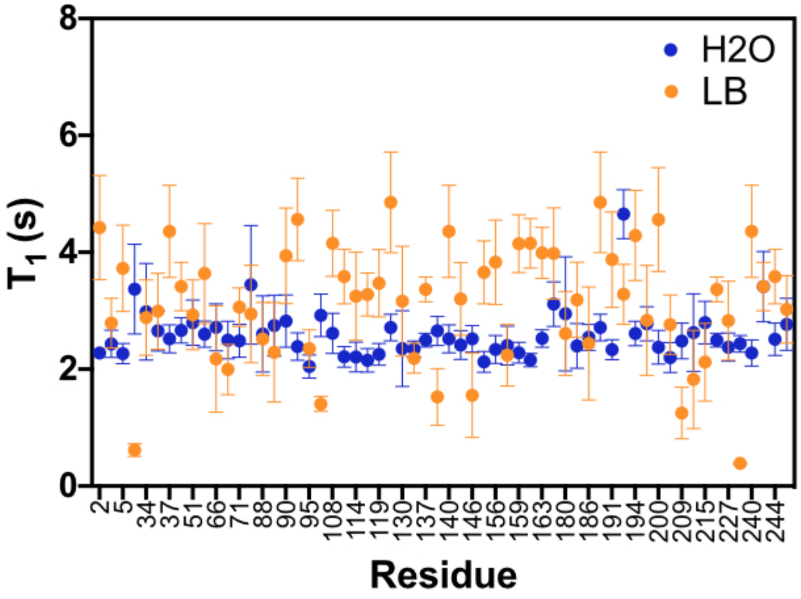


**Figure S5 –** $T_1$ values measured for $^{15}N$ IGPS sample grown in 10% ($H_2O$, $^{14}N$) and 10% LB perdeuterated M9 media. The average $T_1$ values for 10% ($H_2O$, $^{14}N$) sample is 2.6 s ± 0.4, and for 10% LB it is 3.1 s ± 1.

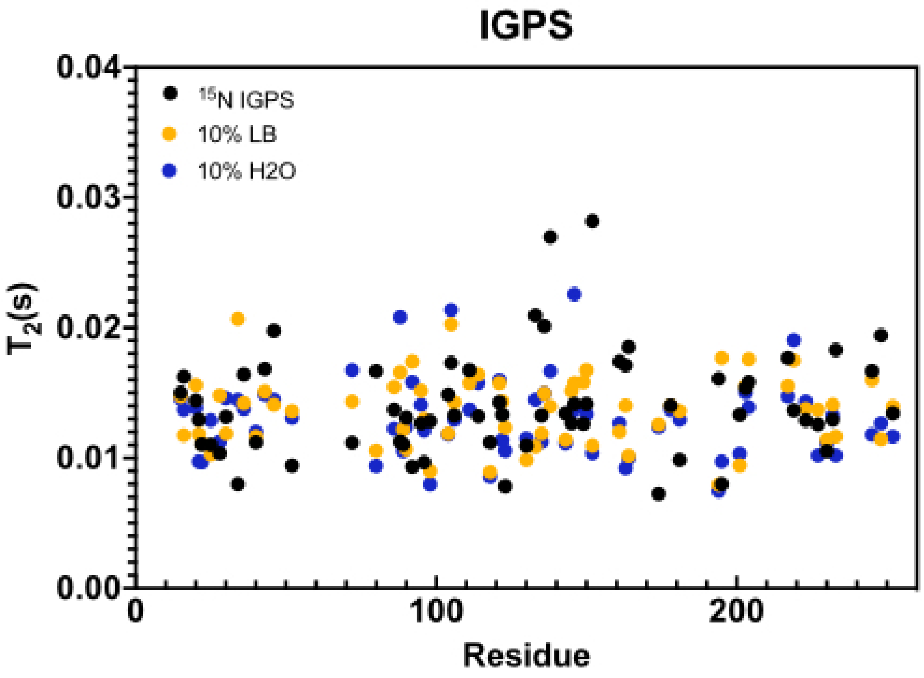


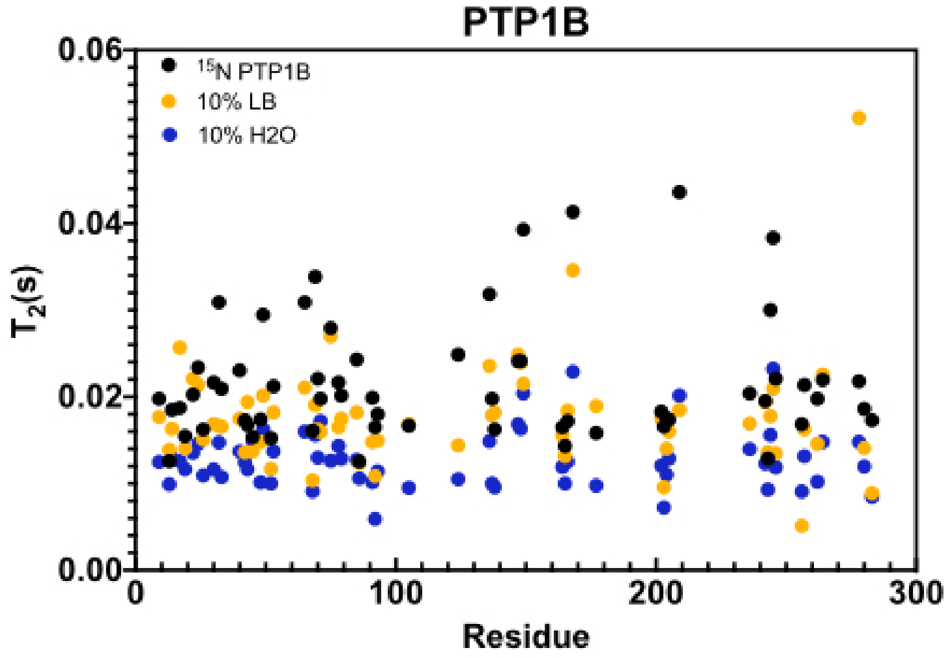


**Figure S6 –** $T_2$ values estimated from NMR resonance peak line width measurements for $^{15}N$ IGPS and PTP1B samples grown in 100% $D_2O$ M9 media, with 10% $H_2O$, and 10% LB addition. Line widths at half height ($\nu_{1/2}$) are determined from integration of NMR resonance peaks with a Lorentzian fit in NMRFAM-Sparky. $T_2$ values are calculated from the following equation: $\nu_{1/2}=1/(\pi \times T_2)$. The average $T_2$ values for fully perdeuterated growth of IGPS and PTP1B are 0.014 ± 0.04 and 0.021 ± 0.007 respectively. For 10% $H_2O$ sample, the IGPS and PTP1B $T_2$ values were determined to be 0.014 s ± 0.003 and 0.013 ± 0.003, and for 10% LB it is 0.013 s ± 0.003 and 0.018 s ± 0.006 respectively.

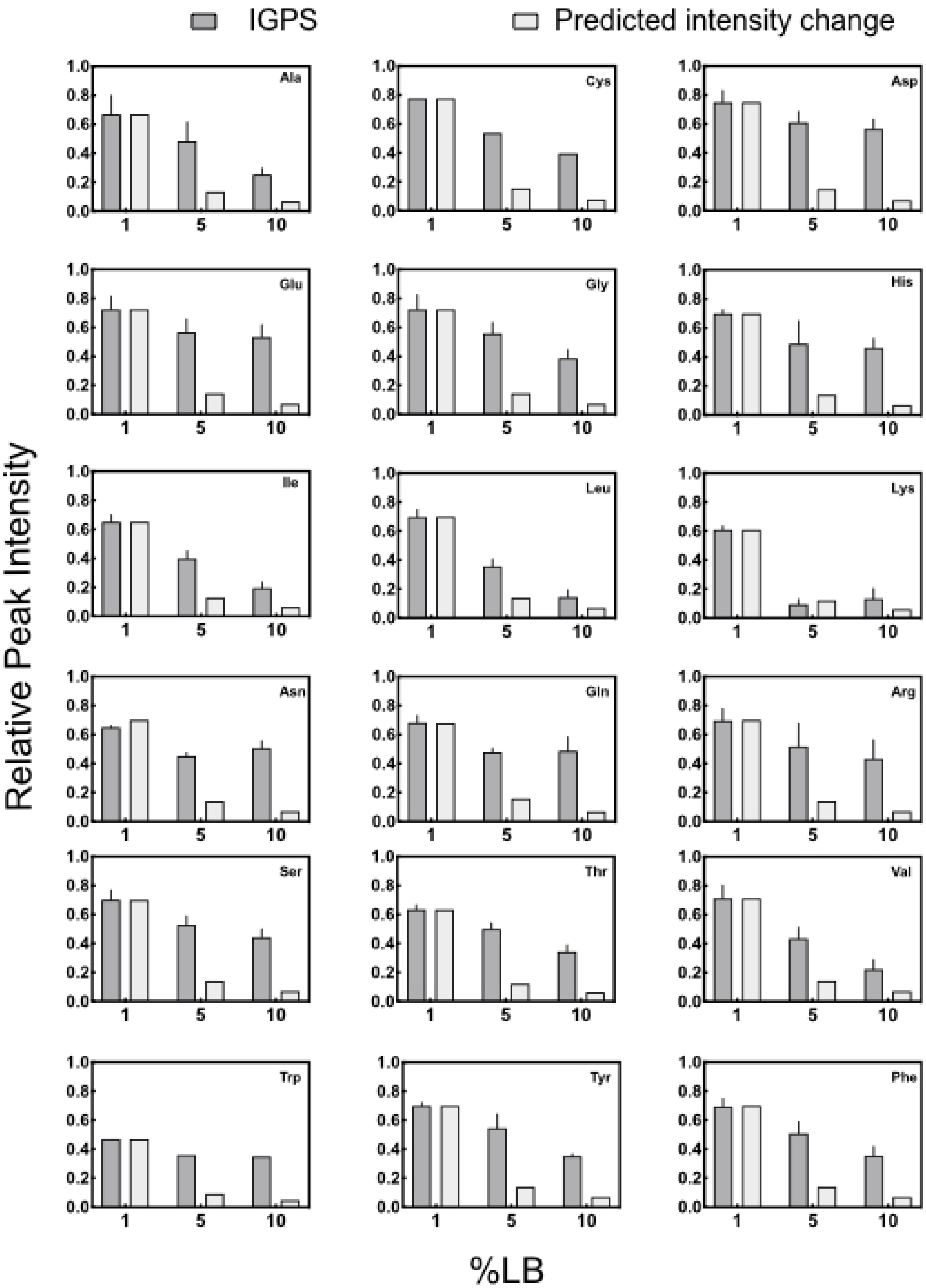


**Figure S7.** Predicted relative intensity changes (light grey) for 5% and 10% LB experiments extrapolated from 1% IGPS data compared to real values obtained from the 5% and 10% LB NMR experiments (dark grey). This analysis reveals that the NMR signal attenuation for Lys is LB concentration dependent. Glu and Asp residues have the highest signal retention. This is because the Glu biosynthetic pathway is responsible for the assimilation of inorganic nitrogen into other amino acid biosynthetic pathways. Additionally, both Glu and Asp are substrates for transamination reactions for all other amino acid biosynthesis pathways (shown in Fig. 1).

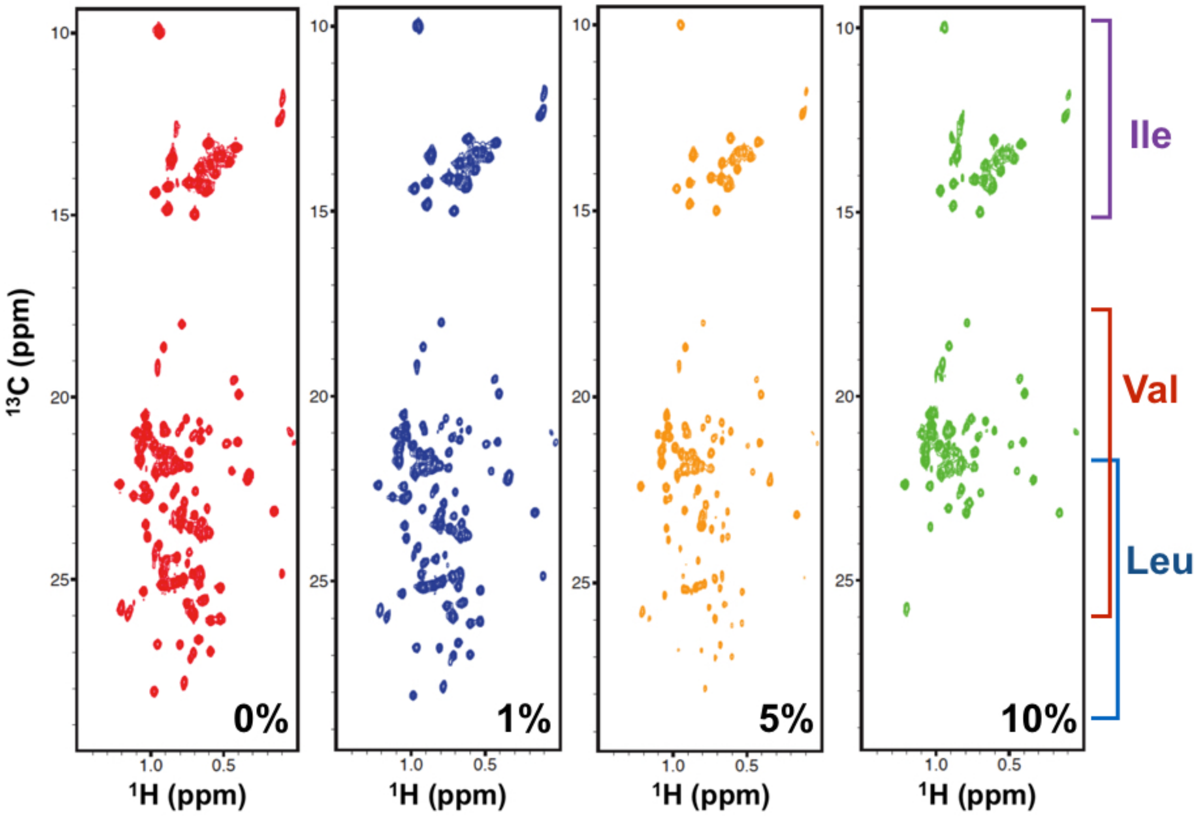


**Figure S8.** Effects of LB doping on ILV spectra. $^{1}H^{13}CH_3$ HMQC NMR spectra of Ile, Leu, and Val methyl groups in IGPS depicting changes in peak intensities in the presence of 1% (blue), 5% (orange), and 10% by volume, LB (green) in the growth medium, relative to that of a reference spectrum (red) without added LB. Leu resonances are strongly affected by LB and disappear gradually from the spectrum and altogether at 10% LB.

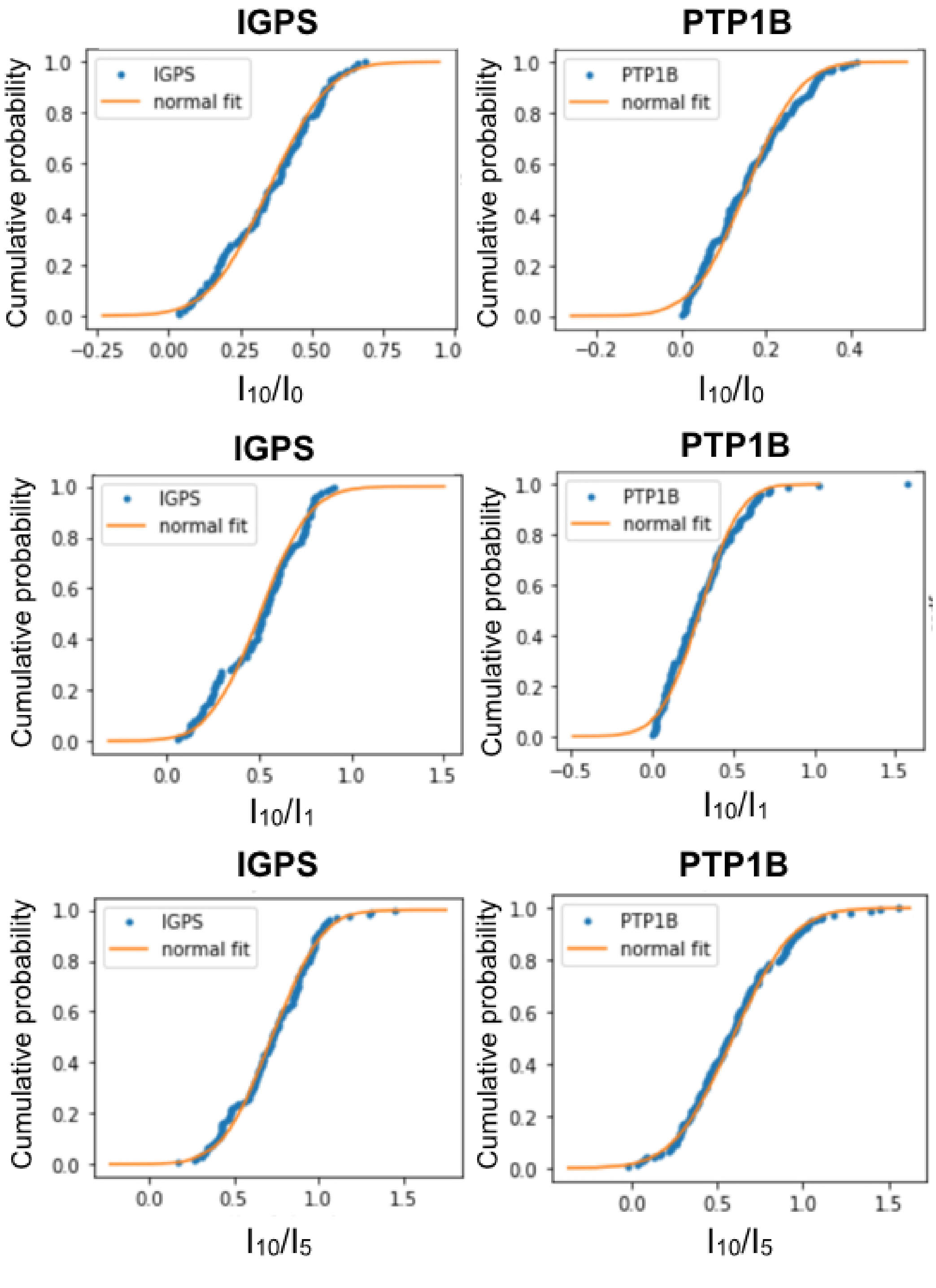


**Figure S9.** The empirical distribution function for signal intensity ratios of 10% LB over 0%, 1% and 5% LB is shown for IGPS and PTP1B. These signal ratios are shown to have a normal distribution.

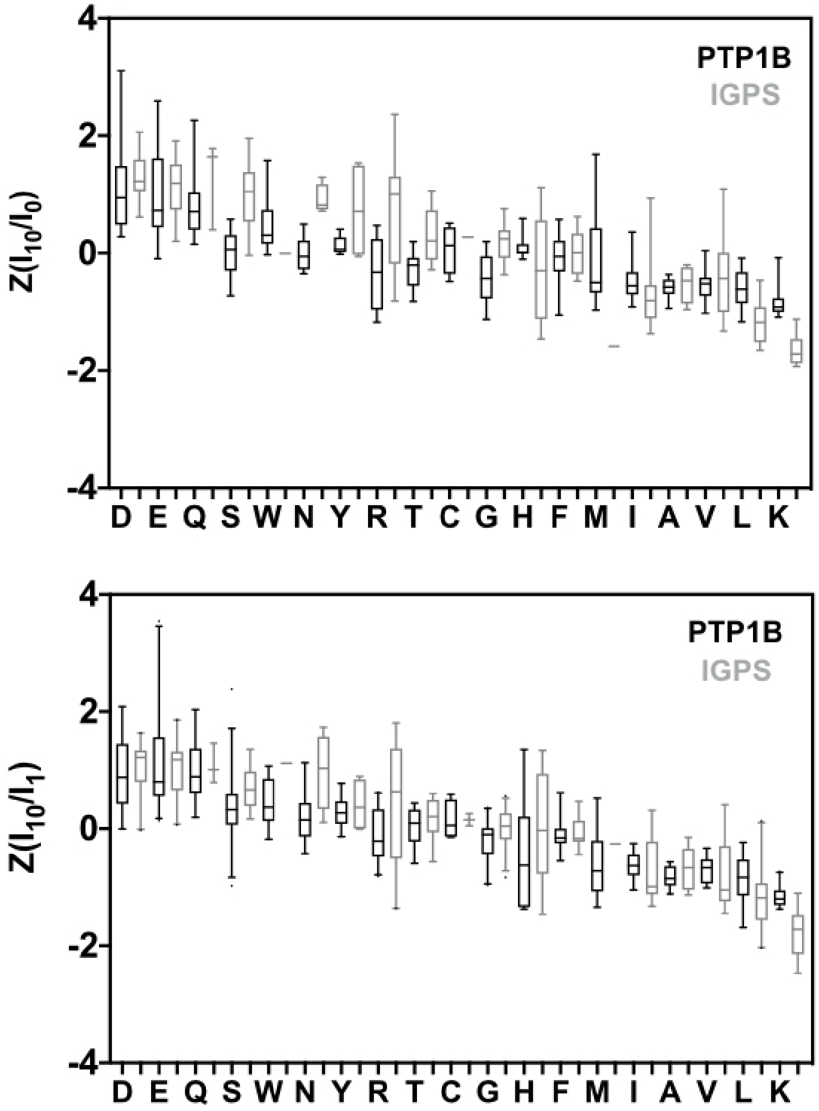


**Figure S10.** Boxplots of $Z(I_{10}/I_0)$ and $Z(I_{10}/I_1)$ values in the order of high to low median values shown for the 8hr and 20hr growths of IGPS and PTP1B. The box represents the interquartile range (25–75 percentile), the median is shown with a line, and the whiskers represent 1-99 percentile range.

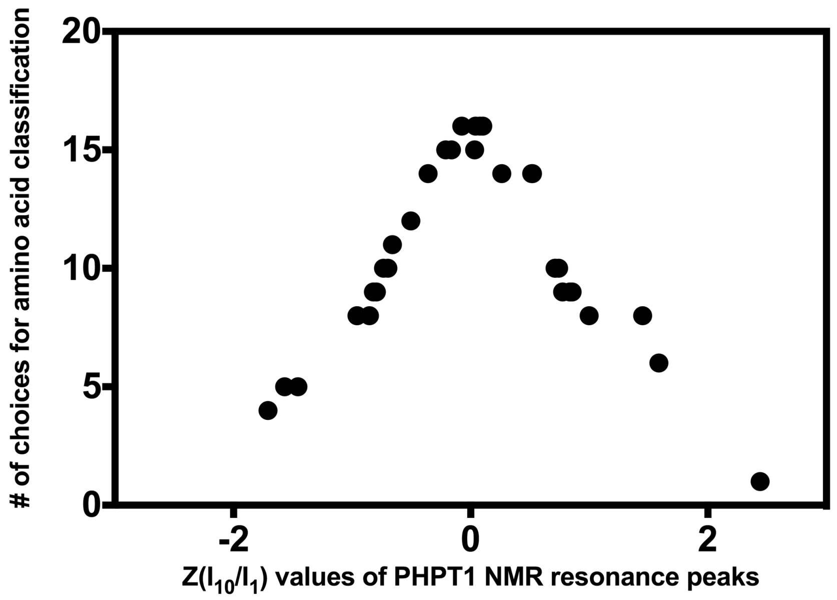


**Figure S11.** The distribution of the number of potential amino acid assignments for residues with $Z(I_{10}/I_1)$ values that are within range of the PHPT1 test data set plotted against the $Z(I_{10}/I_1)$ values of the PHPT1 test data set. The data clearly shows the ambiguity of amino acid type assignment for residues with $-0.7 < Z(I_{10}/I_1) < 0.7$.

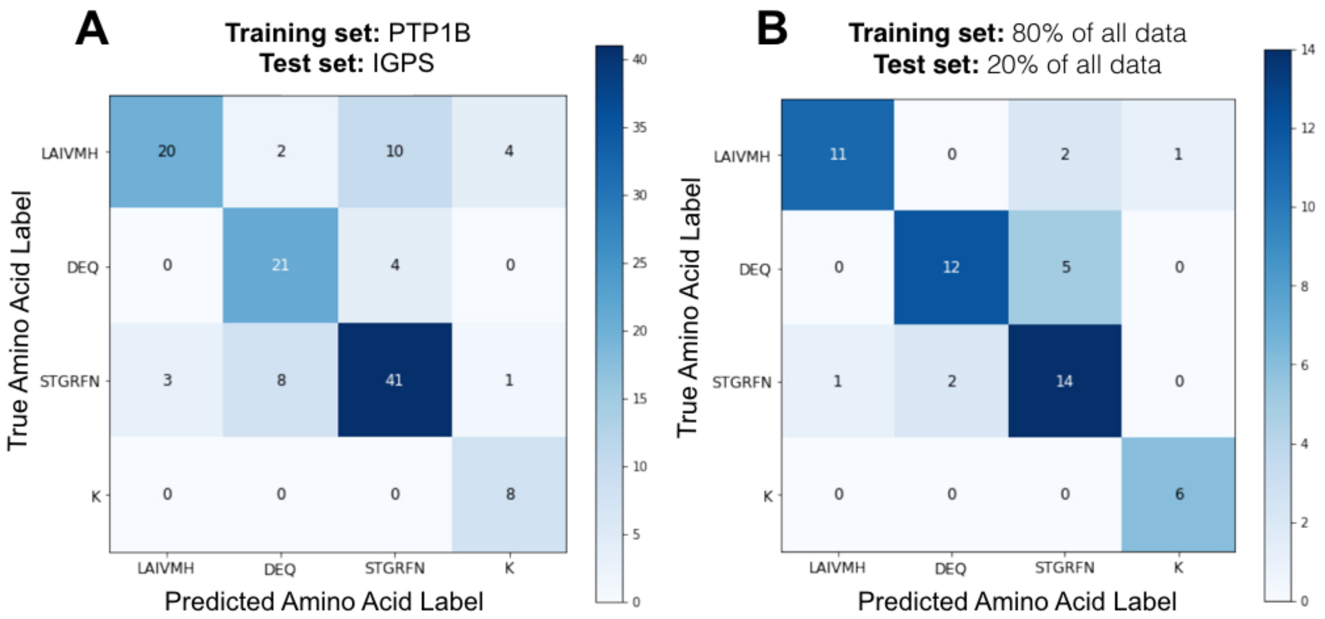


**Figure S12.** Confusion matrices summarizing the prediction power for **A)** a model trained on PTP1B data (n=147) and tested on IGPS data (n=122), where the resulting accuracy was 74%, and **B)** a model trained on combined data in a 80% /20% split of the training and test data. This model had an accuracy of 80%.

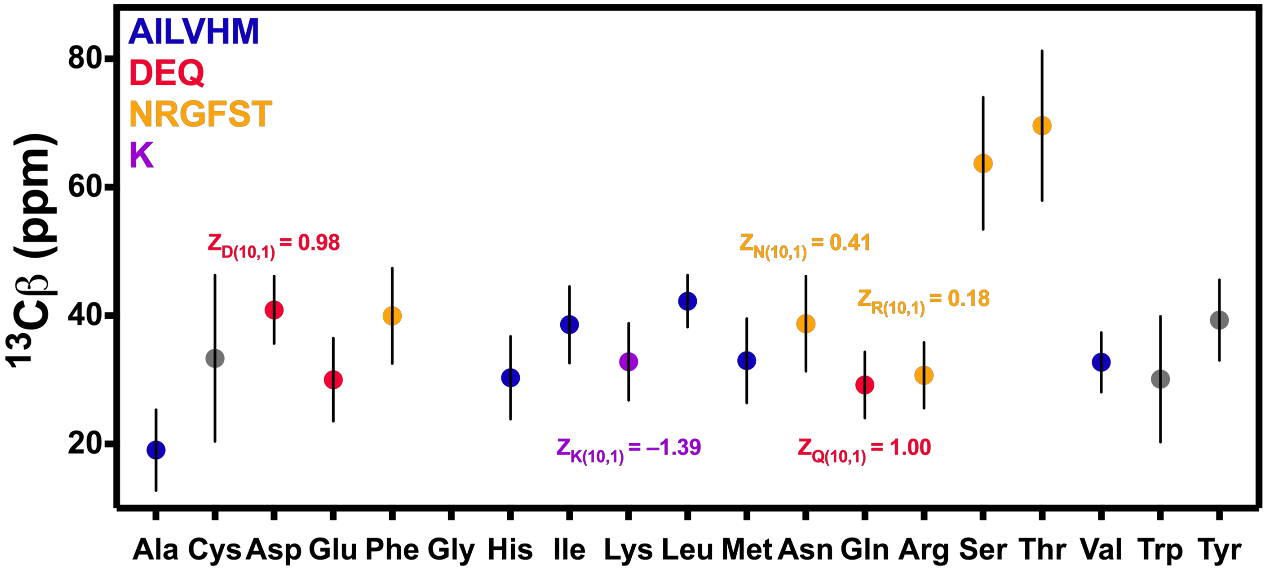


**Figure S13.** The average and standard deviation of $^{13}$Cβ chemical shifts obtained from the Biological Magnetic Resonance Bank (BMRB). LB doping experiments enable the separation of amino acid types with degenerate Cβ chemical shifts. $Z(I_{10}/I_1)$ values for select residues are shown above that amino acid type.

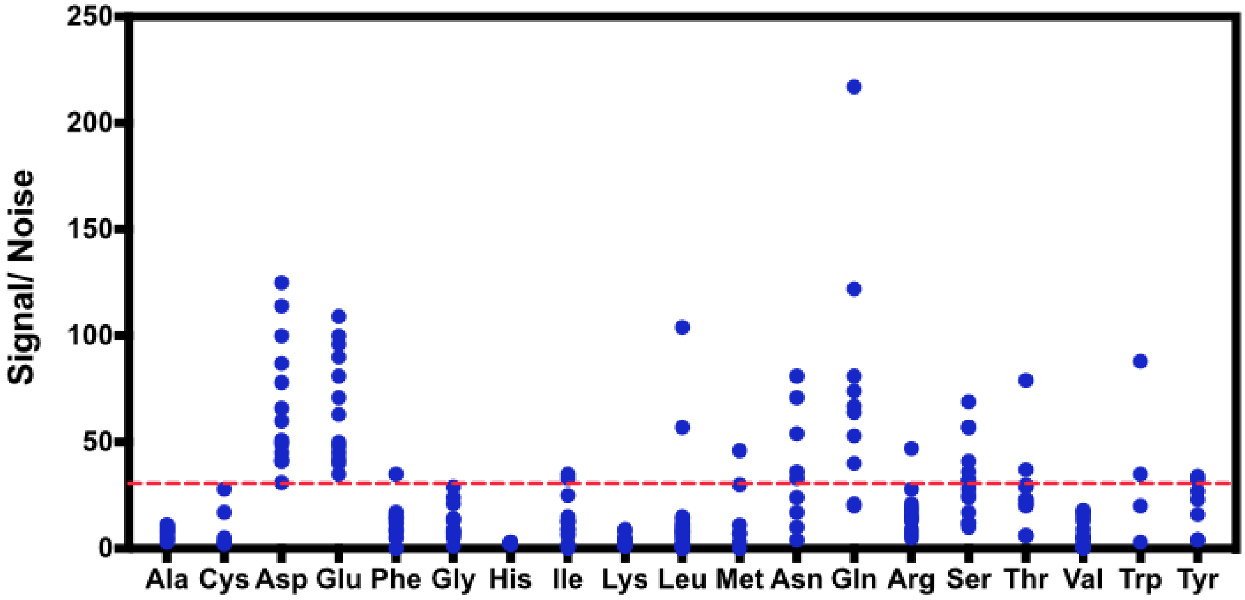


**Figure S14.** The signal to noise of NMR resonance peaks observed in 10% LB $^{15}N$ HSQC spectrum of 336 μM PTP1B. A total of 67 residues are observed with S/N > 30 (indicated with a red line).

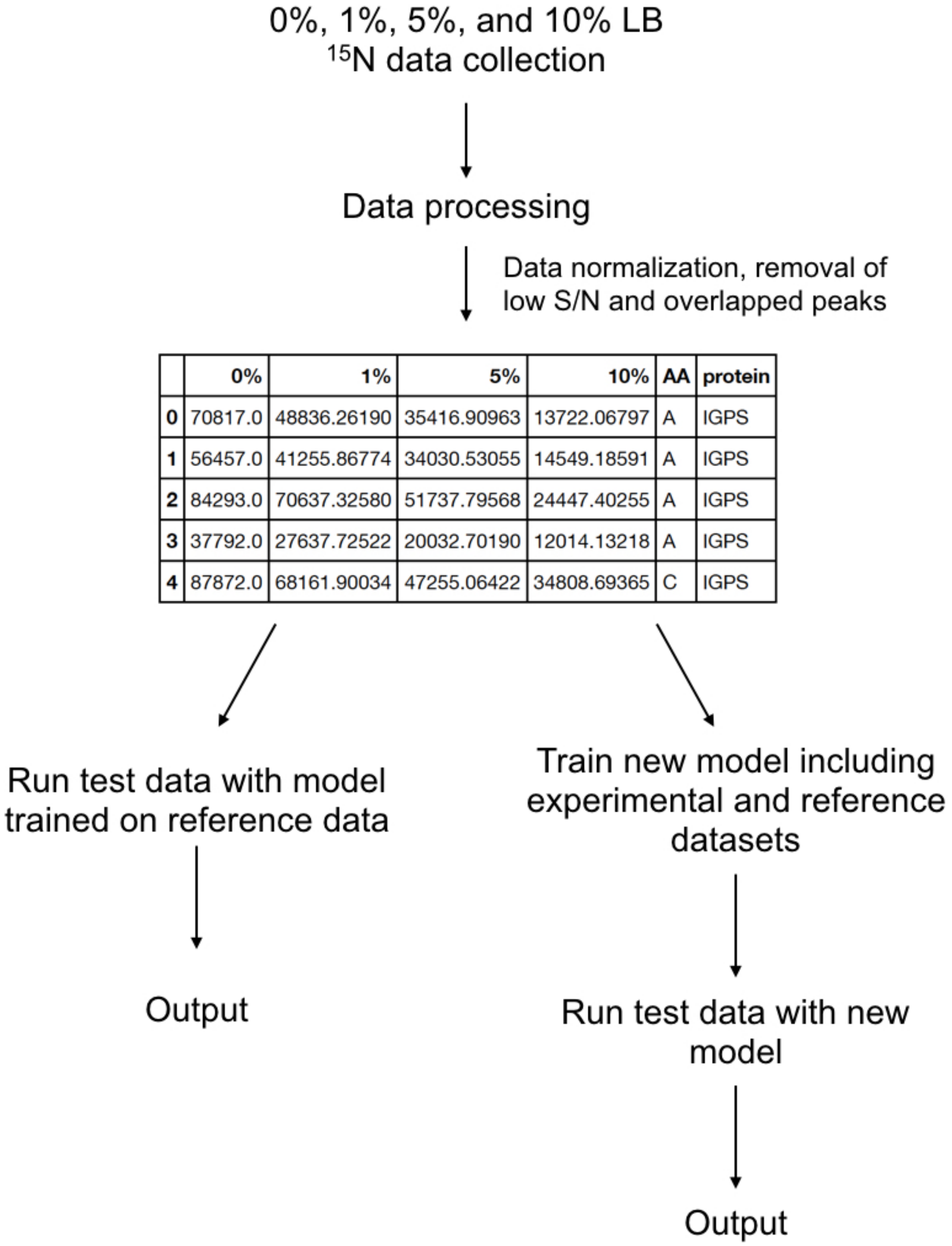


| | 0% | 1% | 5% | 10% | AA | protein |
|---|---|---|---|---|---|---|
| 0 | 70817.0 | 48836.26190 | 35416.90963 | 13722.06797 | A | IGPS |
| 1 | 56457.0 | 41255.86774 | 34030.53055 | 14549.18591 | A | IGPS |
| 2 | 84293.0 | 70637.32580 | 51737.79568 | 24447.40255 | A | IGPS |
| 3 | 37792.0 | 27637.72522 | 20032.70190 | 12014.13218 | A | IGPS |
| 4 | 87872.0 | 68161.90034 | 47255.06422 | 34808.69365 | C | IGPS |

**Figure S15.** Suggested workflow for usage of MeTA prediction tool. It is recommended that all NMR experiments be conducted with the same parameters. The model optimization documentation is available at github (https://github.com/evan-anderson/MeTA) .

**Table S1.** $Z(I_{10}/I_1)$ values of PHPT1 test data (n=34) . Protein was expressed at 25ºC for 16 hr.

| | $Z(I_{10}/I_1)$ | | $Z(I_{10}/I_1)$ |
|---|---|---|---|
| **A50** | -0.795 | **R45** | 0.521 |
| **A54** | -0.959 | **R78** | 0.515 |
| **C73** | -0.211 | **S61** | 0.714 |
| **D58** | 2.298 | **S80** | 0.837 |
| **E51** | 1.450 | **S94** | 0.743 |
| **E72** | 1.587 | **T106** | 0.711 |
| **G62** | 0.078 | **V23** | -0.6o61 |
| **G75** | 0.262 | **V27** | -0.853 |
| **G77** | 0.043 | **V44** | -0.820 |
| **G92** | 0.102 | **V90** | -0.958 |
| **G98** | -0.076 | **Y47** | 0.999 |
| **H102** | -0.735 | **K110** | -1.569 |
| **I43** | -0.359 | **V118** | -0.698 |
| **K41** | -1.709 | **T119** | 0.775 |
| **K48** | -1.458 | **N122** | 0.856 |
| **L6** | -0.504 | **G124** | -0.162 |
| **R26** | 0.033 | **L74** | -0.359 |

### SI Methods:

#### IGPS $^{13}CH_3$ ILV labeling experiment

Separate samples of IGPS were produced in which isotopic labeling of isoleucine, leucine, and valine (ILV) methyl groups was achieved by adding 60 mg/L of alpha-ketobutyric acid [methyl-$^{13}$C;3,3-D2] and 100 mg/L of alpha-ketoisovaleric acid [3-methyl-$^{13}$C;3,4,4,4-D2] (Cambridge Isotope Labs) 30 minutes prior to induction.(Tugarinov et al. 2006) Cells were allowed to reach an $OD_{600}$ of 0.8 – 1.0 before induction with 1 mM IPTG. Cells were expressed under two post-induction conditions: 8 hrs at 37 °C, and 20 hrs at 20ºC. The cells (containing isotopically labeled HisF and perdeuterated HisH) were harvested by centrifugation and resuspended in a buffer of 10 mM Tris, 10 mM CAPS, 300 mM NaCl and 1 mM β-mercaptoethanol at pH 7.5 and then co-lysed by ultrasonication. The cell lysate was clarified by centrifugation and the IGPS complex was purified by Ni-NTA affinity chromatography utilizing the C-terminal histidine tag on HisH. IGPS samples for NMR study were concentrated to 0.38 – 0.41 mM in a buffer containing 10 mM HEPES, 10 mM KCl, 0.5 mM EDTA, and 1 mM DTT at pH 7.3.

Experiments probing the side chain methyl groups of Ile, Leu, and Val ($^{13}CH_3$-ILV) were based on the multiple-quantum (MQ) pulse sequence.(Mueller 1979) In these experiments, transmitter offsets alternated between the water resonance and the center of the methyl region (0.75 ppm), while the $^{13}$C channel offset was set to 19.5 ppm. All $^{1}H^{13}CH_3$-ILV spectra were collected with 32 transients, 144 $t_1$ increments, 3778 data points, and spectral widths of 8500 Hz (direct) and 3500 Hz (indirect). NMR spectra were processed with NMRPipe (Delaglio et al. 1995) and analyzed in PINE-SPARKY (Lee et al. 2009). NMR resonance signal intensities in 0%LB PTP1B were multiplied by two to account for the collected transient differences between experiments.
